# The changing landscape of VREfm in Victoria, Australia: a State-wide genomic snapshot

**DOI:** 10.1101/289975

**Authors:** Robyn S. Lee, Anders Gonçalves da Silva, Sarah L. Baines, Janet Strachan, Susan Ballard, Glen P. Carter, Jason C. Kwong, Mark Schultz, Dieter Bulach, Torsten Seemann, Timothy P. Stinear, Benjamin P. Howden

## Abstract

Vancomycin-resistant *Enterococcus faecium* (VREfm) represent a major source of nosocomial infection worldwide. In Australia, the *vanB* genotype is dominant; however there has been a recent increase in the predominantly plasmid-encoded *vanA* genotype, prompting investigation into the genomic epidemiology of VREfm in this context.

**Materials and Methods:** A cross-sectional study of VREfm in Victoria, Australia (Nov.10^th^ - Dec.9^th^, 2015). A total of 321 VREfm isolates (from 286 patients) were collected and whole-genome sequenced with Illumina NextSeq. Single nucleotide polymorphisms (SNPs) were used to assess relatedness. Multi-locus sequence types (STs), and genes associated with resistance and virulence were identified. The *vanA*-harbouring plasmid from an isolate from each ST was assembled using long-read data.

**Results:** *vanA*-VREfm comprised 17.8% of isolates. ST203, ST80 and a *pstS*(-) clade, ST1421, predominated (30.5%, 30.5% and 37.2% of *vanA*-VREfm, respectively). Most *vanB-*VREfm were ST796 (77.7%). *vanA*-VREfm isolates were closely-related within hospitals vs. between them (core SNPs 10 [interquartile range 1-357] vs. 356 [179-416] respectively), suggesting discrete introductions of *vanA*-VREfm, with subsequent intra-hospital transmission. In contrast, *vanB*-VREfm had similar core SNP distributions within vs. between hospitals, due to widespread dissemination of ST796. Overall, *vanA*-harbouring plasmids differed across STs, and with exception of ST78 and ST796, Tn*1546* transposons also varied.

**Conclusions:** *vanA*-VREfm in Victoria is associated with multiple STs, and is not solely mediated by a single shared plasmid/Tn*1546* transposon; clonal transmission appears to play an important role, predominantly within, rather than between, hospitals. In contrast, *vanB-*VREfm appears to be well-established and widespread across Victorian healthcare institutions.

## Introduction

*Enterococcus faecium* is a leading cause of nosocomial infections world-wide (1, 2). Resistance to glycopeptides, especially vancomycin, is the most clinically-relevant for *E. faecium* (1); mortality associated with any *E. faecium* blood-stream infection (BSI) exceeds 30% (3), while patients with vancomycin-resistant *E. faecium* (VREfm) bacteremia have ∼2.5-fold higher odds of death compared to those with vancomycin-susceptible infection (4). Resistance to vancomycin in *E. faecium* can be conferred by different *van* gene clusters (*A, B, D, E, G, L, M,* and *N* (5–8)), with most infection attributed to *vanA* and *vanB* genotypes (9). VREfm is thought to be transmitted between hospitalized patients, healthcare staff, and/or via the hospital environment (10), with risk of colonization highest among those receiving long courses of antibiotics, the critically-ill and/or immunosuppressed, and those with prolonged hospitalization or history of nursing home residence (11). VREfm has also been shown to arise *de novo* in a patient, following transfer of *vanB* gene cluster from commensal gut anaerobes to *E. faecium* (12, 13).

The prevalence of the *vanA* and *vanB* genotypes varies geographically; in North America, *vanA* has been shown to drive the VREfm epidemic (e.g., (14)), while in Europe, both *vanA* and *vanB* genotypes play a role (15). In Australia, the first case of VREfm detected in 1994 carried the *vanA* operon (16); however, it is *vanB*-VREfm that has subsequently predominated.

Multiple clones of *vanB*-VREfm have been detected in Australia, mostly from sequence types (STs) within the hospital-associated clonal complex (CC) 17(9, 17–20). A survey of sentinel laboratories from across Australia in 2011 found that 97.2% (104/107 isolates) of vancomycin-nonsusceptible isolates had the *vanB* genotype (17), over half of which were ST203 (19, 21). Subsequent country-wide surveys showed this strain dominated until 2014, when it was surpassed by the ST796 clone (22) - largely due to its widespread dissemination throughout Victoria and southeastern Australia (23).

Along with the spread of the *vanB-*associated ST796 clone, there has been a recent, dramatic increase in *vanA*-VREfm; since 2011, the prevalence of *vanA*-VREfm bacteremia, as measured by these sentinel surveys, increased from <1% (2/341) to 21% (90/408 isolates genotyped) in 2016 (24). The reasons for this increase remain unclear, but multiple STs appear to be involved. A retrospective study examining *vanA*-VREfm (screening and clinical isolates) across four Australian hospitals from 2011-2013 (25) suggested this increase could not be attributed to a single ST or *vanA*-harbouring plasmid; however, this was based on a convenience sample of only 18 isolates from across the whole country, making this inconclusive.

To further explore the potential reasons for the shift in *van* genotype, and increase our understanding of the molecular epidemiology of VREfm, we conducted a population-level study of VREfm in the State of Victoria – the second most inhabited State in Australia (http://www.abs.gov.au/AUSSTATS/). All VREfm isolates from colonization and clinical infection detected across the State over a month were subjected to whole genome sequencing, as well as isolates from VSEfm bacteremia. In doing so, we provide an in-depth assessment of the genomic diversity of *E. faecium* and novel insights into the epidemiology of VREfm in this region.

## Materials and Methods

### Study design and population

A cross-sectional survey of VREfm was conducted between November 10^th^ – December 9^th^, 2015 in the State of Victoria (population: 5,931,100 for 2015, per the Australian Bureau of Statistics; http://www.abs.gov.au/AUSSTATS/). During this period, all VREfm-positive isolates (including screening and clinical samples) from all laboratories across the State were sent to the Microbiological Diagnostic Unit Public Health Laboratory (MDU), as well as all vancomycin-susceptible *E. faecium* isolated from blood cultures. Final lists of isolates received by the MDU were cross-checked with primary diagnostic laboratories to ensure all eligible isolates were included.

### DNA extraction and WGS

Genomic DNA was isolated from a single colony using a JANUS Chemagic Workstation and Chemagic Viral DNA/RNA kit (PerkinElmer). Libraries were prepared with the Nextera XT DNA sample preparation kit (Illumina). Whole genome sequencing (WGS) was performed using the Illumina NextSeq 500 with 150 base-pair paired-end reads.

### Bioinformatics

Sequences were analysed using the Nullarbor pipeline (Seemann T, available at: https://github.com/tseemann/nullarbor).

In brief, Illumina reads were trimmed for quality using Trimmomatic (v.0.36, (26)). Contamination was assessed using Kraken (v.0.10.5, (27)). Reads were assembled into contigs using SPAdes (v.3.10.1, (28)). *In silico* identification of resistance genes was done with ABRicate (Seemann T, available at: https://github.com/tseemann/abricate), wherein the assembled contigs were searched for resistance genes in the ResFinder database using BLAST+ (29). Putative virulence genes were also identified in the same manner using the Virulence Factor DataBase (30), to assess whether increased virulence could be responsible for the rise in VREfm bacteremia in Australia. All positive hits were reviewed for gene completeness (i.e., gene coverage and gaps), percent identity, and depth of coverage. BLAST+ (29) was also used to search the assembled contigs for *E. faecium* alleles listed in the pubMLST database (https://pubmlst.org); unknown allelic combinations and allelic variants were submitted to the database for new assignment. Single nucleotide polymorphism (SNP)-based analyses were done by aligning reads using the Burrows-Wheeler Aligner MEM algorithm (v.0.7.16, (31)) to a novel local Victorian reference (ST796, AUSMDU00004028; see **Table A1** for details and metrics). This reference was selected because preliminary analyses indicated the majority of our samples were also ST796. A minimum mapping quality of 60 was required, and ambiguously-mapped reads were excluded. SNPs were called using Freebayes (v1.0.2, (32)) under a haploid model, with a minimum depth of coverage of 10 and allelic frequency of 0.9 required to call a SNP. Integrative Genomics Viewer (v.2.3.88) was used to review alignments and SNPs as needed (33).

### Phylogenetics and population structure

ClonalFrameML was used to identify potential recombination (v.1.11, (34)). Putative recombination sites were then masked using a custom script (available at: https://github.com/kwongj/cfml-maskrc), and concatenated core SNP sites were extracted using snp-sites -c (35). A maximum likelihood tree was then produced using RAxML (v.8.2.11, (36)) under a General Time Reversible model with four gamma categories. One thousand bootstrap replicates were performed to assess confidence in the tree. Figures were produced using ggtree (37) in R (v.3.3.2). Bayesian Analysis of Population Structure with hierarchical model-based clustering (hierBAPS; v.6.0, (38)) was also used to identify major clades, using the alignments of recombination-adjusted core SNPs as input.

### Comparative genomics of vanA-VREfm

One strain from each of the major *vanA*-harbouring STs was sequenced using PacBio Single Molecule, Real-Time sequencing with P6-C4 chemistry (Pacific Biosciences, CA). *De novo* assembly was done using Canu (v.1.3 (39)); see **Appendix** for further detail. QUAST (v.4.5) was used for additional quality metrics (40). *vanA*-harbouring plasmids from each of these strains were identified using ABRicate, and subsequently annotated using Prokka (v. 1.11, (41)) with the *Enterococcus* database (https://github.com/tseemann/prokka/blob/master/db/genus/Enterococcus). To compare the gene content of *vanA*-harbouring plasmids, these plasmids were aligned to one another using progressiveMauve (build date 2014-12-19) and visualized in Geneious (v.9.1.7, (42)). Tn*1546* transposons were identified and genes were manually compared across plasmids.

### Victorian isolates in global context

To assess how Victorian *E. faecium* strains fit in the global context, we re-ran our analyses including 877 additional strains from Europe (872), North America (2), Africa (2) and Asia (Israel; 1) (43–46), and isolates from the Welcome Trust Sanger BSAC Resistance Surveillance Project (from NCBI BioProject PRJEB344). A maximum likelihood tree was produced using IQ-TREE (v.1.6.1, (47); see **Appendix**) with model selection based on the lowest Bayesian Information Criterion.

### Statistical analysis

Pairwise SNP distributions were compared using the Mann-Whitney rank-sum test. Differences in proportion were compared using either the Chi-Square test or Fisher’s Exact test, as appropriate. Analyses were done in Stata (v.14.2, College Station, TX: StataCorp).

### Data availability

Illumina data, and the PacBio reads with corresponding assemblies for the five isolates listed in **Table A1** are available on the National Center for Biotechnology Information’s (NCBI) Sequence Read Archive under BioProject PRJNA433676.

## Results

Between November 10^th^ – December 9^th^ 2015, 333 *E. faecium*-positive samples were detected at 19 primary diagnostic laboratories from across Victoria (two additional laboratories reported no *E. faecium* during this time). Two eligible VSEfm isolates were not submitted, thus 331 isolates were available for sequencing. All isolates passed WGS quality control (see the **Appendix** for details).

In total, 321 isolates (from 286 patients) were VREfm - 20 of which were from BSI (from 17 patients). Ten VSEfm isolates (from ten patients) were also collected; one of these was likely a false negative, as the patient also had a *vanA*-VREfm isolated from blood collected the same day (further detail in the **Appendix**). Thus, 27 patients had *E. faecium* bacteremia over this period; an incidence of 5.5/100,000 when extrapolated to one year.

Most patients (n=257, 87.1%) were from hospitals in major metropolitan centers, compared to rural areas (n=29, 9.8%), consistent with the geographical distribution of the population in this State. Overall, 35 hospital networks (HCNs) were represented, with isolates received from up to five affiliated hospitals from a single network. Five isolates were collected in community medical/dental clinics (3.1%). Sex and age were not available for patients from HCN1 (n=61, 20.7%); among the remaining patients, 113 were male (48.3%), and the median age was 71 (range 18-98, IQR 58-80). At sample collection, 256 persons were inpatients and 36 were outpatients. Four patients were missing location of isolation and had unknown admission status.

Twenty-seven patients had >1 sample positive for *E. faecium* (range 2-5 per patient; **Table A2**), with a maximum of 40 SNPs between repeat isolates of the same ST (**Figure A1**). Repeat samples were collected within the same hospital as the original sample for 26/27 patients (96.3%, **Table A2**).

### Identification of van genotype

*In silico* genotyping revealed that 260 (78.6%) of isolates were exclusively *vanB*, and 59 (17.8%) were exclusively *vanA*. There was no association between *van* genotype and BSI (**Figure 1**, Fisher’s Exact p=0.52). Two isolates were positive for *vanA* and *vanB* (<1%). These were most closely related to *vanB*-VREfm isolates, with as few as 0 pairwise core SNPs compared to *vanB*-VREfm isolates (vs. 195 for *vanA-*VREfm), suggesting they already had *vanB* when *vanA* was acquired. As expected, all isolates with *vanA* and/or *vanB* were phenotypically resistant to vancomycin (MIC > 4 mg/L (48)). Neither *vanA* or *vanB* were found in isolates classified as VSEfm (3.0%).

**Figure 1.**
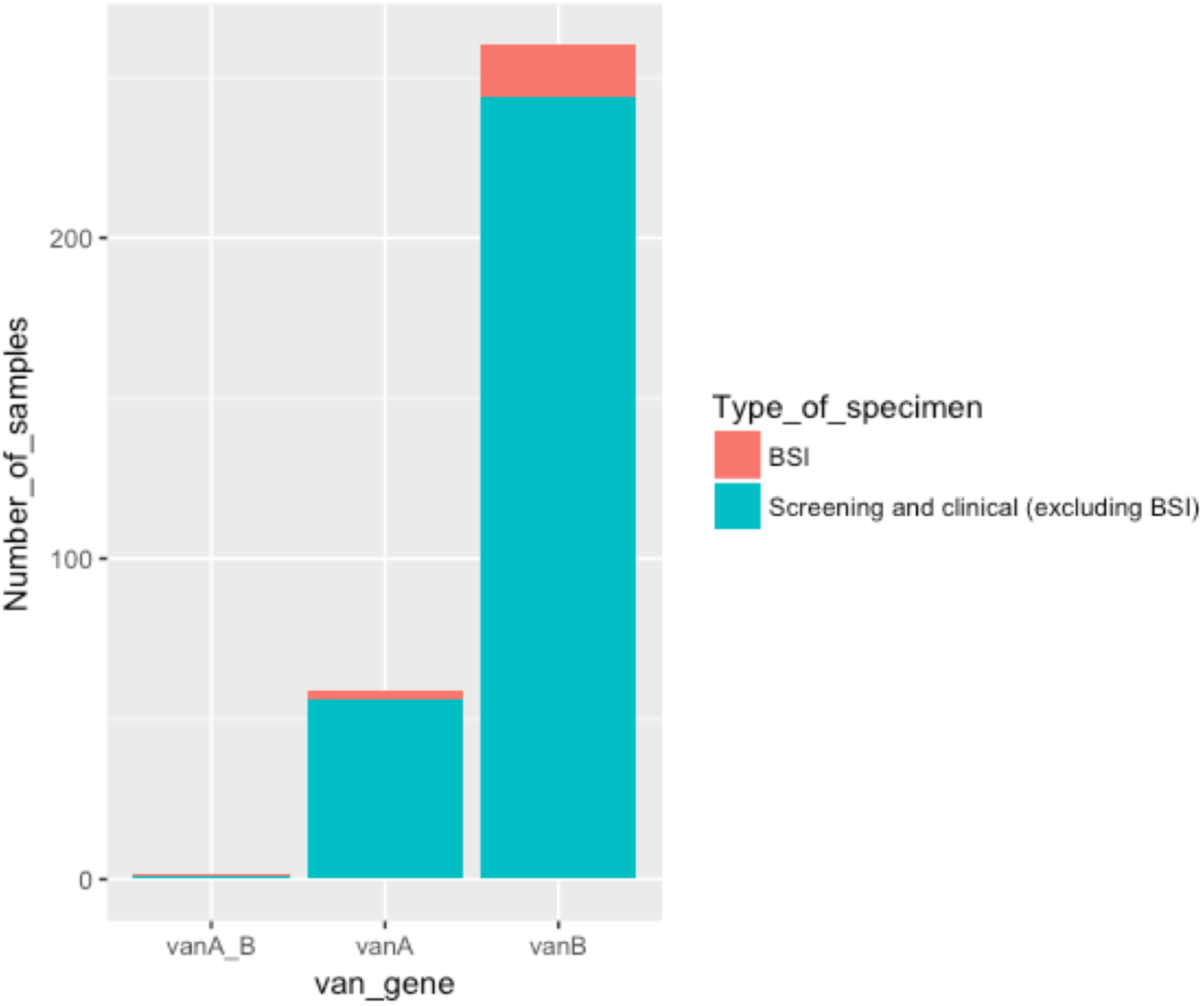
Specimen type by *van* gene for Victorian *E. faecium* isolates. Only 2 isolates were *vanA+vanB;* only one was from a blood-stream infection (BSI). As all 10 VSEfm isolates were from BSI by design, these have not been shown. Fisher Exact test comparing the proportion of BSI among isolates with *vanA* vs *vanB*: p= 0.52.

Overall, *vanB*-VREfm was more widely dispersed across Victorian hospitals and HCNs compared to *vanA*-VREfm; excluding the two *vanA+vanB-*positive samples, *vanB*-VREfm isolates were identified in 34/35 HCNs, while *vanA*-VREfm was only identified in 10/35 during this time period (p < 0.00005). VSEfm isolates were from six different networks.

### Multi-locus sequence typing

Eighteen previously-known MLSTs and four novel STs were identified. Most isolates were ST796 (61.3%, **Table 1**). The *pstS* allele was missing for 22 (6.6%) isolates; these were classified as ST1421 based on remaining alleles (**Table A3**).

**Table 1.**
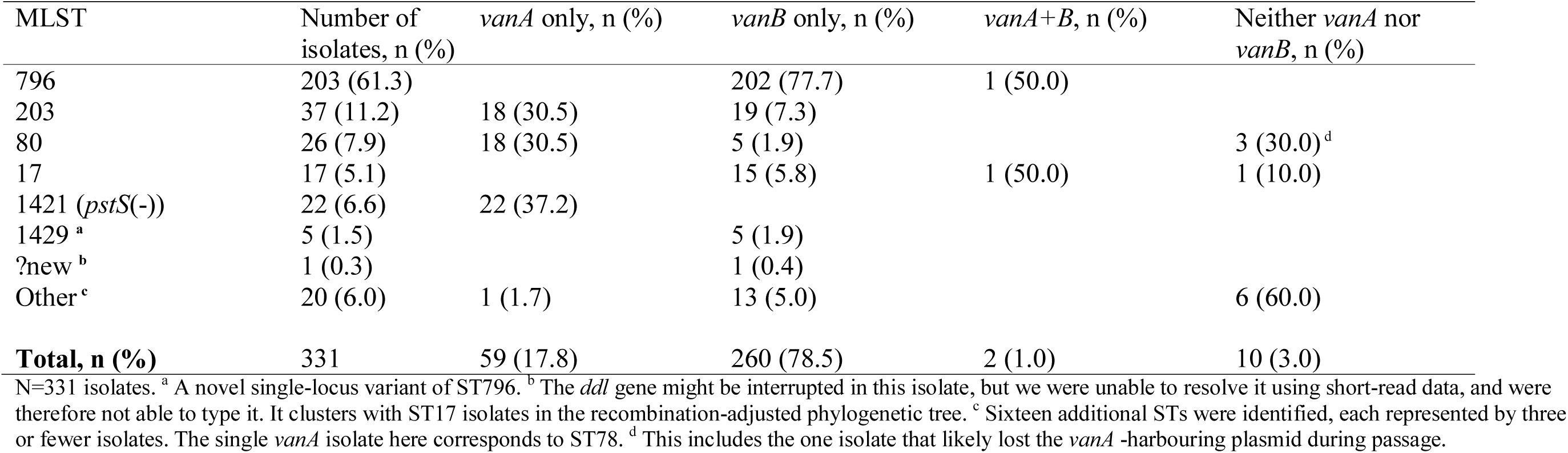
Multi-locus sequence types (MLSTs) identified.

*vanA* was found in fewer STs than *vanB* (3 vs. 11, respectively, excluding *vanA+B* positive isolates), though this trend was not statistically significant (Fisher’s Exact test, p=0.17). *vanA* was almost exclusively present in ST203 and ST80, and ST1421 isolates (**Table 1**). Most *vanB*-positive isolates were ST796.

### Phylogenetics and population structure

Prior to adjusting for recombination, there were a median of 20 core SNPs between study isolates and the reference (IQR 14-2,653); adjusting for recombination reduced this to a median of 12 core SNPs (IQR 9-238). An alignment of these adjusted core SNPs was used to produce a maximum likelihood tree (**Figure 2**; recombination blocks are shown in **Figure A2**). Many branches had very low bootstrap support, therefore BAPS was used to validate clusters.

**Figure 2.**
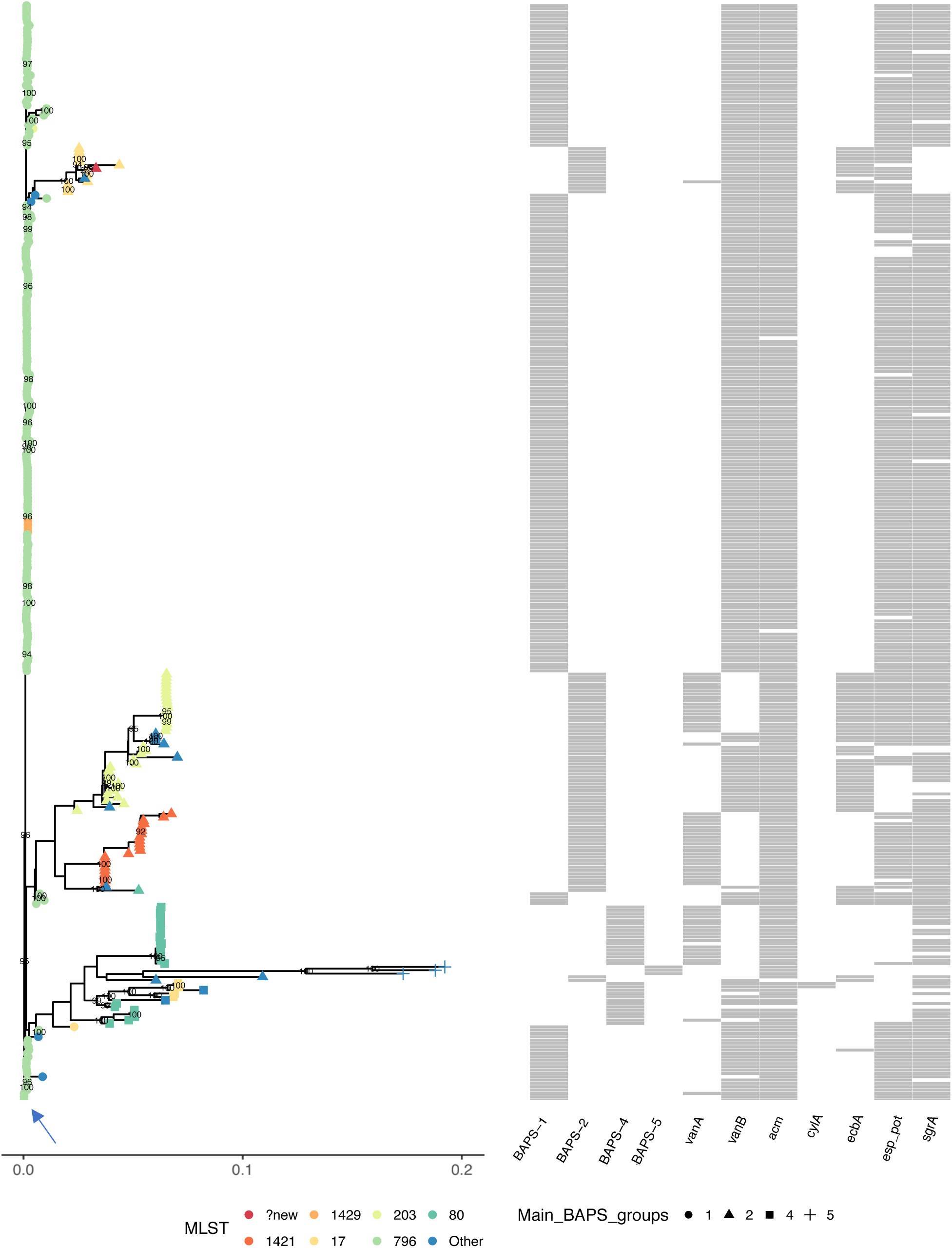
Maximum likelihood tree of *E. faecium* in Victoria. 6,497 recombination-adjusted core SNPs from all 331 isolates (321 VREfm and 10 VSEfm) were concatenated, and used to produce a maximum likelihood tree in RAxML (36) under a General Time Reversible model with gamma rate heterogeneity. One thousand bootstrap replicates were performed to assess confidence in the phylogeny. BAPS-3 is not shown; AUSMDU00004157 (an ST54 VSEfm) was the only isolate in this clade, and was >3,800 SNPs from all other isolates, so it was excluded from this figure for easier visualization. Branches with bootstrap support >90 are shown. Many branches had very low bootstrap support, thus hierarchical Bayesian Analysis of Population Structure (hierBAPS) was used to further validate the clusters identified. Main BAPS groups are shown. The reference is identified with an arrow. No isolates had *ebpA*, *hyl*, *gelE*, *agg, pilA* or *pilB,* therefore these genes have not been shown.

Five major BAPS groups were identified, which were largely consistent with the phylogeny (**Figure 2**). BAPS-1 was comprised predominantly of ST796 isolates (203/214, 94.9%), and was highly clonal, with a median of 10 core SNPs separating isolates (IQR 7-15). In contrast, BAPS-2 was comprised of multiple STs. The highest proportions were from ST203 (36/82, 43.9%), ST1421 (22/82, 26.8%) and ST17 (12/82, 14.6%). Isolates in BAPS-2 were separated by a median of 257 core SNPs (IQR 167-312). Excluding repeat samples from the same patient, 81 pairs of isolates were within one SNP. Sixty-six of these pairs (81%) were isolated at the same hospitals as one another (H1, H3, H5), while the remainder were predominantly from different HCNs (see **Figure A3**). BAPS-3 was comprised of a single VSEfm isolate from ST54, which was over 3,900 SNPs from other isolates (not shown in **Figure 1**). BAPS-4 isolates were predominantly from ST80 (25/31, 80.6%), but also included isolates from ST17, ST78 and ST262. Isolates were separated by a median of 252 core SNPs (IQR 6-346). All pairs within one core SNP were from ST80; excluding repeat isolates, 8/15 of these pairs were collected at the same hospitals as one another (H1, H2 and H4). BAPS-5 was comprised of three VSEfm isolates (ST21, ST22, and ST32), with a minimum of 369 core pairwise SNPs.

### Genetic diversity across hospitals and van genotypes

Excluding comparisons within the same patient (shown in **Figure A2**), the median pairwise core SNP distances among VREfm were similar within and between hospitals, at 196 (IQR 13-297) and 193 (IQR 13-207), respectively (**Figure 3A**).

**Figure 3.**
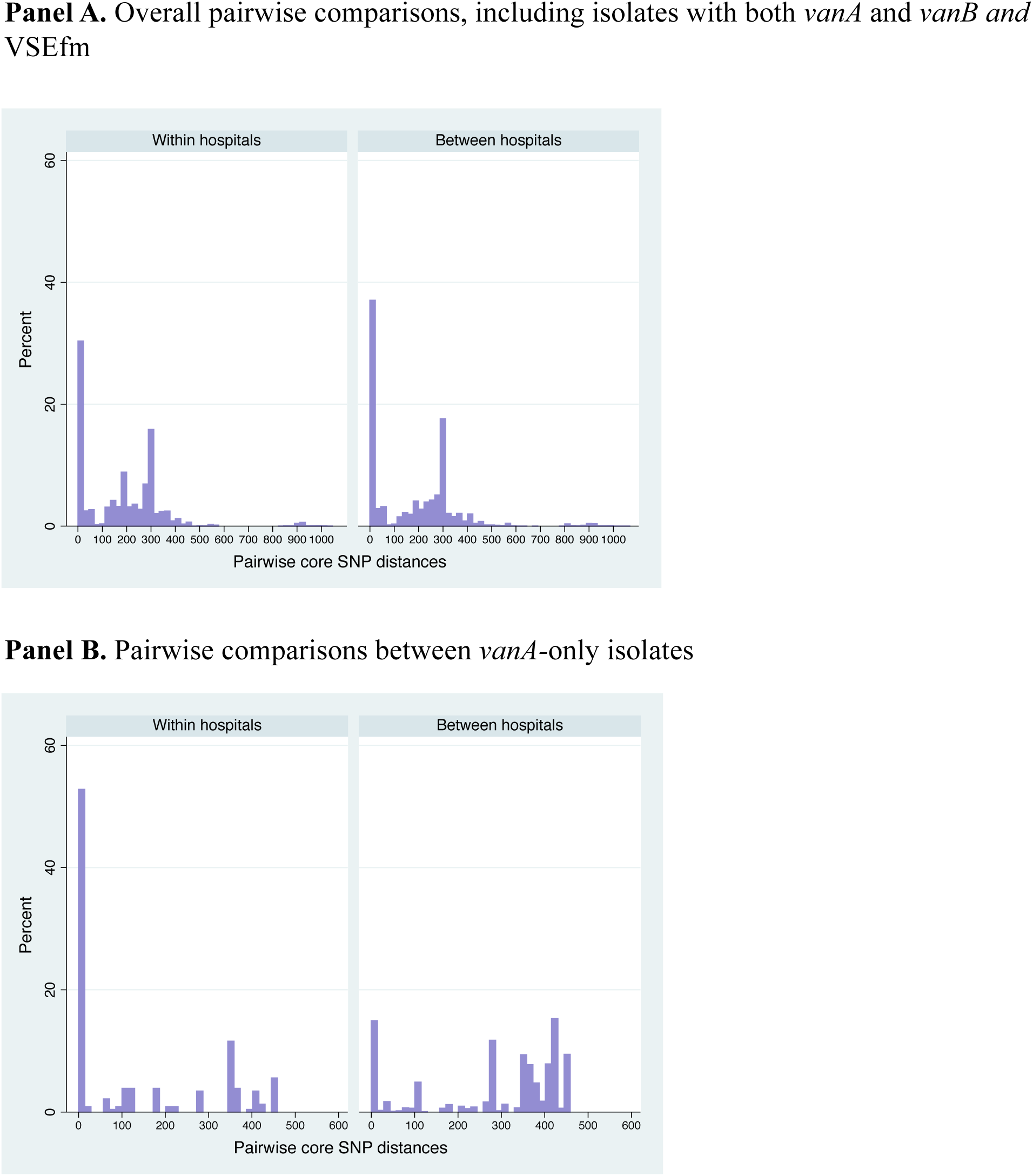
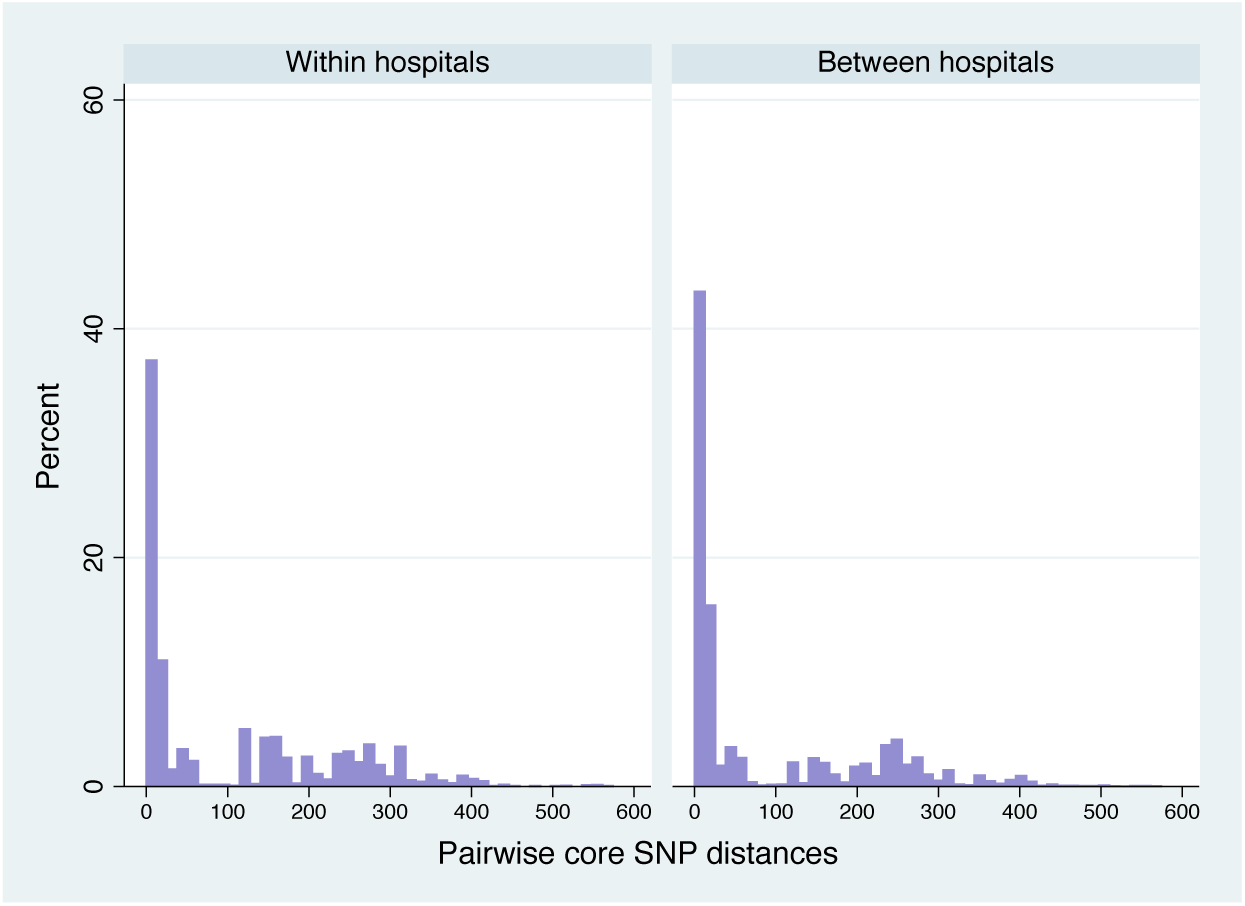
Recombination-adjusted pairwise core SNPs separating Victorian isolates. **A.** Pairwise SNP comparisons are shown regardless of *van* gene. There were 54,567 total pairwise comparisons, after excluding those involving isolates with missing data on hospital/HCN, those from unknown clinics, and pairs within the same patients. Between hospitals, the median pairwise SNPs was 193 (IQR 11-298), while within hospitals, the median was 196 (IQR 13-297). For easier visualization, as one VSEfm isolate (AUSMDU00004157) was a minimum of 3,893 SNPs from all others, only comparisons between the 330 other isolates are shown in the figure. **B.** Pairwise comparisons between isolates with *vanA* only are shown. There were 1,706 such comparisons (233 within hospitals 1,473 between hospitals). The median pairwise SNPS within hospitals was 10 (IQR 1-357) compared to a median of 356 (IQR 179-416) between hospitals (p=0.0001). **C.** Pairwise comparisons between isolates with *vanB* only are shown. There were 33,639 such comparisons (3,104 within hospitals, 30,528 between hospitals). The median pairwise SNPs within hospitals was 40 (IQR 9-206), compared to a median of 15 (IQR 8-174) between hospitals (p=0.0001). Isolates with both *vanA* and *vanB*-positive (n=2), and isolates that were *vanA* and *vanB*-negative (n=10) were excluded from Panels B and C.

As we hypothesized there were differences in diversity between isolates with different VREfm genotypes, *vanA-*VREfm and *vanB*-VREfm were analyzed separately. *vanA*-VREfm were significantly less diverse within hospitals compared to between them (**Figure 3B**, p=0.0001), suggesting multiple, independent introduction events with subsequent intra-hospital transmission. In contrast, *vanB*-VREfm was more diverse within hospitals vs. between them (**Figure 3C**), p=0.0001), consistent with wide-spread dissemination and long-term establishment of these strains across institutions within the Victorian healthcare system.

### Virulence

Isolates were interrogated for the presence of putative virulence factors; as shown in **Figure 2**, only two isolates (both ST17) had the *cylA* gene which is required for expression of cytolysin (49). No genes encoding hyaluronidase (*hyl*_Efm_ (50)), agglutination substance (*agg/asa1* (51) or gelatinase (*gelE* (52)) were detected; the latter two genes are thought to be more prevalent in *E. faecalis* than *E. faecium* (53).

Microbial surface components recognizing adhesive matrix molecules including *acm* (54), which has been found predominantly in clinical isolates, as well as *ecbA* (55) and *sgrA* (55) were also investigated. *acm* was present in 98.5% of study samples, across nearly all STs. *ecbA* was present in 62 isolates (30.5% of *vanA-*VREfm; 15.8% of *vanB*-VREfm), including all ST192, ST400 and ST893 isolates, and most ST17 and ST203 isolates (71% and 97%, respectively). *sgrA* was present in 294 isolates (91.9%, of *vanA*-VREfm; 89.2% of *vanB*-VREfm) from nearly all STs. No isolates had *pilA* and *pilB*, which are also potentially involved in adhesion (56).

Finally, the gene *esp,* which has previously been associated with hospital outbreaks (57) and encodes enterococcal surface protein (58), was investigated. While repeat regions made it impossible to completely assemble *esp* using short-read data (see **Appendix, Figure A4**), we found that 266 isolates had at least 80% coverage (81.6% of *vanA*-VREfm; 86.2% of *vanB*-VREfm), suggesting that these isolates were likely *esp*-positive. In this context, *esp* was more prevalent in bacteremia due to VREfm than VSEfm (17/20 vs. 1/9, p=0.0001, excluding the VSEfm that had likely lost the *vanA* operon).

The *vanA*-harbouring plasmid from one isolate per ST was completely assembled using long-read data; no putative virulence genes were found to co-localize with *vanA* on the same plasmid (**Figure A5**).

### Comparative genomics of vanA-VREfm

The *vanA*-harbouring plasmids from ST78 and ST796 had the greatest homology (**Figure 4**), with similar plasmid backbones and 100% amino acid identity of the Tn*1546* transposon, which carries the *vanA* gene cluster (**Figure A5**). The main difference between these plasmids was a large insertion in ST796 compared to ST78, carrying *repB,* IS*1216* and two hypothetical proteins (**Figure A5**). The *vanA*-harbouring plasmid from ST80 was similar to that from ST796 (**Figure A6**), but had lost ISEfa7-*birA* from the plasmid backbone (**Figure A5**) and had a 48-bp deletion in *vanS* compared to the ST796 plasmid. Both *vanA*-harbouring plasmids from the ST203 and ST1421 isolates had unique plasmid backbones and Tn1546 transposons (**Figure A5**). Together, these findings suggest that horizontal transfer of a single *vanA*-harbouring plasmid and/or Tn*1546* carrying the *vanA* gene cluster is not responsible for the increase in *vanA*-VREfm in Victoria.

**Figure 4.**
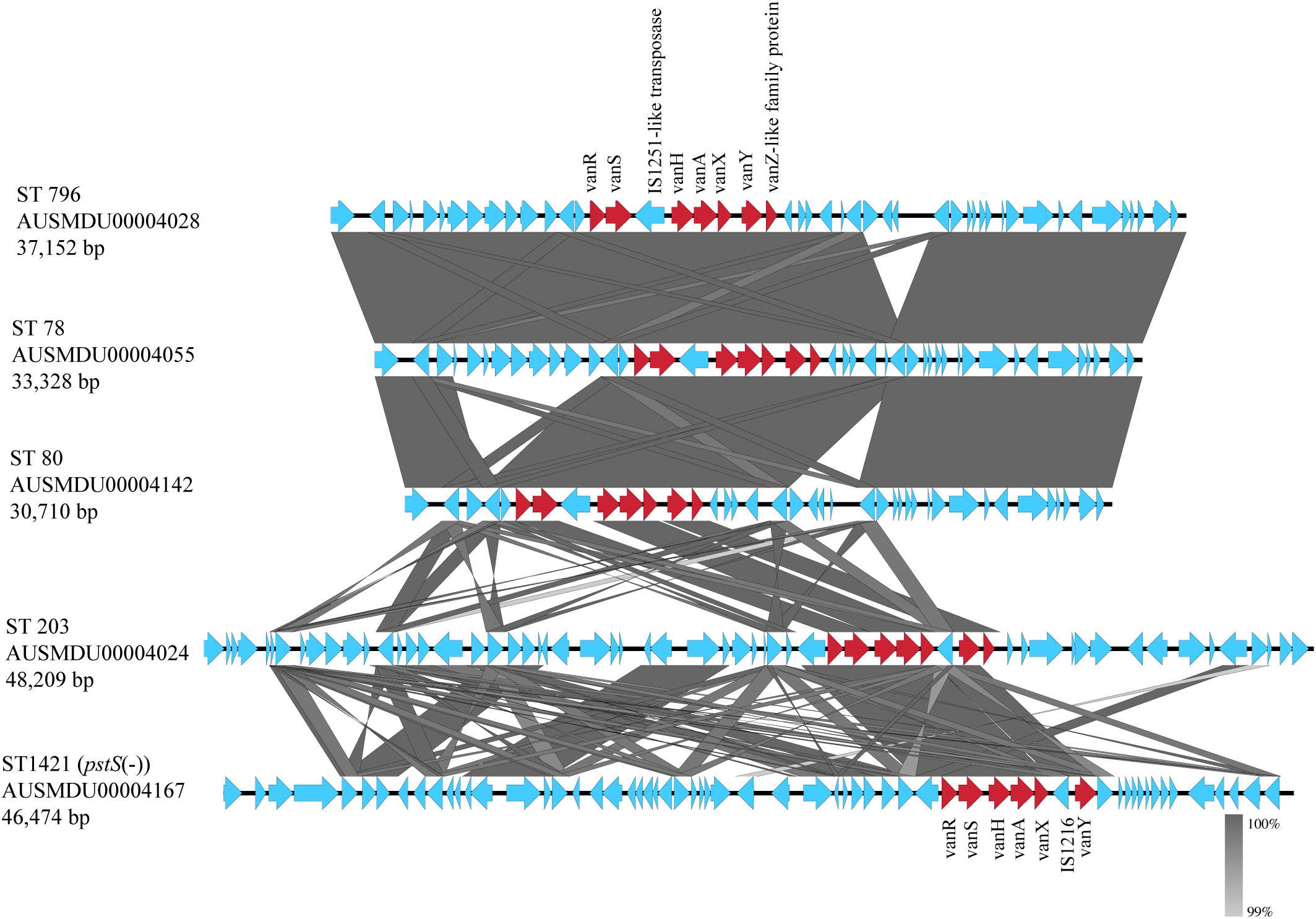
Homology between *vanA*-harbouring plasmids. PacBio Single Molecule Real-Time sequencing was performed on a *vanA-*positive isolate from each of the main *vanA*-associated STs (ST 78, ST80, 203, 796, and ST1421). Plasmids harbouring *vanA* were initially identified using ABRicate (https://github.com/tseemann/abricate), and subsequently annotated using the *Enterococcus* database (https://github.com/tseemann/prokka/blob/master/db/genus/Enterococcus). This figure was produced using EasyFig (v. 2.2.2, version); BLASTn was used to compare sequence homology. Only hits with a minimum length of 200 bp and at least 99% identity are shown for clarity. *van* genes are indicated in red, all other genes are indicated in blue.

### Victorian isolates in global context

Compared to published sequences, the *vanA*-harbouring plasmids from ST78, ST796 and ST80 shared closest homology to a *vanA*-harbouring plasmid from VREfm in Demark (46) (**Table A4**). The *vanA*-harbouring plasmid from ST203 was most closely related to plasmids from several STs of VREfm isolated in Western Australia and Queensland (25), while the plasmid from ST1421 was closely-related to that from another ST1421 *E. faecium,* isolated in New South Wales in 2014 (59) (**Table A4**).

Phylogenetic analysis based on core SNPs from Victorian isolates and global *E. faecium* strains showed that, with the exception of those from ST796, Victorian isolates were widely distributed across the tree (**Figure A7**). This suggests ongoing, inter-continental transmission of *E. faecium*.

## Discussion

For the past two decades, the majority of VREfm in Australia have been *vanB* genotype (13, 18, 19, 21–23, 60–62). Recently, however, *vanA*-VREfm has been increasing. In this population-based study, we looked at factors potentially contributing to this change in epidemiology.

Firstly, we examined the genetic diversity of *vanA*-VREfm across our State, to assess for potential transmission. Consistent with the hypothesis of van Hal et al. (25), we found that a single clone is not driving this increase; instead, our findings suggest that multiple introductions of *vanA*-VREfm have occurred, with subsequent spread within Victorian hospitals. This pattern is similar to that of early *vanB*-VREfm in Australia (63), though the latter has become dominated by a single ST in Victoria (ST796) (23).

Another possibility was that a single *vanA*-harbouring plasmid and/or Tn*1546* transposon was being transmitted within and across different STs, as has been shown in Denmark (46). As a unique *vanA*-harbouring plasmid was present in representatives from each ST, and nearly all had different Tn*1546* transposons, this suggests that horizontal gene transfer is not the sole driver of *vanA*-VREfm in Victoria.

Finally, we assessed whether there were differences in putative virulence factors between our isolates, and those from other studies/settings. Overall, few putative virulence factors were identified, and their prevalence was largely consistent with previous data from *vanB*-VREfm in Australia (64), and similar to or lower than those found in *vanA*-VREfm from settings (e.g., in China (65) or Brazil (66)). While much is still unknown about virulence factors in VREfm (compared to *Enterococcus faecalis*), this suggests the rise in *vanA*-VREfm in Australia, and the continued success of *vanB*-VREfm, is not due to changes in virulence. Consistent with this, the rate of *E. faecium* BSI in our study was similar to that of another population-based study in Denmark (3). This has important implications for hospital infection control; as the patient population affected by VREfm has also remained consistent over time, we propose that environmental factors may be mediating the observed changes in VREfm epidemiology. Further studies are needed to investigate this hypothesis.

In addition to gaining new insights in *vanA* epidemiology in this context, we have also provided valuable information on *vanB*-VREfm; given that we found genetic diversity was slightly lower across hospitals compared to within, this suggests a potential community reservoir for these strains. Transmission of VREfm has previously been shown in nursing homes (67), with carriage strains closely related to those causing infection; investigation is warranted to determine if this is also the case for Victoria. The seemingly widespread diversity of *vanB* VREfm could also be due to a “diffusion” effect, wherein colonization and infection have slowly spread across hospitals through many years of patient exchanges. Such a pattern would be reminiscent of that observed in Denmark (68); however, while in Denmark the “diffusion” process was marked by a hub-and-spoke pattern, with patients mainly being exchanged between regional hospitals and central hospitals in Copenhagen, in Victoria, the process was likely more random. If such a “diffusion” scenario is correct, we hypothesize that the pattern of genetic diversity of *vanA*-VREfm will eventually mirror that of the *vanB*.

This work has a number of key strengths. Firstly, by including >99% of VREfm collected in Victoria over this one-month period, we have provided a comprehensive assessment of local strain diversity and the prevalence of clinically-relevant *van* genotypes, as well as a baseline to monitor for changes in VREfm population in Victoria. Secondly, we used local reference genomes for all SNP-based analyses. As using more genetically-distant reference genomes can result in false positive SNPs, and/or loss of information due to differences in genes present in the study sample, but not in the reference, this increased the accuracy of our short-read analyses. Since the first complete *E. faecium* genome was published by our group in 2012 (69), few other genomes have been completed; by sharing these complete VREfm genomes (from five different STs), we are also providing an invaluable resource for public health as well as future studies on *E. faecium*. Finally, the use of long-read data has also allowed us to completely assemble and characterize the *vanA*-harbouring plasmids in each of the different STs – a task that is not always feasible with short-read data alone.

There are several limitations of this work. Firstly, screening protocols for VREfm colonization are not standardized across Victoria; each hospital adhered to its own infection control guidelines, and as such, there may be incomplete capture of colonizing strains in this dataset. This may bias our dataset somewhat towards strains causing clinical infection, and cause us to miss potential transmission. Another limitation is that we did not have any data on patient contact or intra-hospital transfer (except where isolates were collected at different hospitals) – preventing us from confirming direct person-to-person transmission events. However, we do not think this affects our overall population-level inferences, given the dramatically lower genetic diversity of *vanA* within hospitals compared to *vanB*. Finally, as this was predominantly a lab-based study, we also did not have data on clinical presentation at time of sample collection and were therefore unable to discriminate clinical infection aside from bacteremia. Thus, we may have underestimated the overall prevalence of infection (though the most clinically important cases - associated with bacteremia - have been included).

Herein, we have provided a comprehensive snapshot of VREfm strain diversity across Victoria. We highlight the complexity of *E. faecium* genomic epidemiology, revealing key differences in transmission dynamics of clinically-relevant *van* genotypes in this setting. Our findings are applicable to not only to Victoria, but other regions with VREfm - particularly those where the prevalence of *vanB* is high. While *vanB*-VRE continues to predominate in Victoria, repeated introductions and dissemination of *vanA-*VREfm within our hospitals suggest this is changing. To better understand the clinical significance of this shift, prospective studies with detailed corresponding clinical data are needed. As plasmids carrying *vanA* or their Tn*1546* transposons may be may be readily transferred to VSEfm and/or rarely to other species (e.g., (70)), preventing an increase in *vanA*-VREfm is critical. This reinforces the overall importance of ongoing prevention and control of VREfm.

## Transparency

Authors have no conflicts of interest to declare.

## Funding

RSL is supported by a Fellowship from the Canadian Institutes of Health Research (Funding Reference Number 152448). SLB holds an Australian Government Research Training Program Scholarship. JCK is supported by an early career fellowship from the NHMRC (GNT1142613). BPH has a Practitioner Fellowship from the National Health and Medical Research Council (NHMRC), Australia (GNT1105905), and is supported by the Centre of Research Excellence on Emerging Infectious Diseases (NHMRC GNT1102962). The Microbiological Diagnostic Unit Public Health Laboratory is funded by the Victorian Government, Australia.

## Acknowledgements

We would like to thank the participating laboratories for their provision of samples and the clerical and technical assistance of staff at MDU PHL. We would also like to thank Dr. Sharon Peacock (Department of Medicine, University of Cambridge, Cambridge, United Kingdom; Wellcome Trust Sanger Institute, Hixton, United Kingdom; London School of Hygiene and Tropical Medicine, London, United Kingdom) and Dr. Kathy Raven (Department of Medicine, University of Cambridge, Cambridge, United Kingdom) for helpful discussions about global VREfm.

## Contributions

RSL designed and ran the primary analyses, interpreted results, made the tables and figures, and wrote the first draft of the manuscript. AG designed analyses, advised on bioinformatics approaches, helped interpret results, and contributed to writing the manuscript. SLB assembled the PacBio genomes, provided input on analyses, and helped interpret results. JS recruited all labs, coordinated submission of isolates and data from primary diagnostic laboratories, and did the initial processing of samples at MDU PHL. SB designed sample collection and advised on the initial interpretation of the results. GPC did the laboratory work to generate the PacBio sequences. JCK, MS, DB and TS provided bioinformatics tools, and advised on bioinformatics analyses. TPS and BPH conceived the cross-sectional study, helped decide on analyses, and edited the manuscript. All authors critically reviewed the manuscript for content.

## References

1. Gilmore MS, Lebreton F, van Schaik W. 2013. Genomic transition of enterococci from gut commensals to leading causes of multidrug-resistant hospital infection in the antibiotic era. Curr Opin Microbiol 16:10–6.

2. Guzman Prieto AM, van Schaik W, Rogers MR, Coque TM, Baquero F, Corander J, Willems RJ. 2016. Global emergence and dissemination of enterococci as nosocomial pathogens: attack of the clones? Front Microbiol 7:788.

3. Pinholt M, Ostergaard C, Arpi M, Bruun NE, Schonheyder HC, Gradel KO, Sogaard M, Knudsen JD, Danish Collaborative Bacteraemia. 2014. Incidence, clinical characteristics and 30-day mortality of enterococcal bacteraemia in Denmark 2006-2009: a population-based cohort study. Clin Microbiol Infect 20:145–51.

4. DiazGranados CA, Zimmer SM, Klein M, Jernigan JA. 2005. Comparison of mortality associated with vancomycin-resistant and vancomycin-susceptible enterococcal bloodstream infections: a meta-analysis. Clin Infect Dis 41:327–33.

5. Lebreton F, Depardieu F, Bourdon N, Fines-Guyon M, Berger P, Camiade S, Leclercq R, Courvalin P, Cattoir V. 2011. D-Ala-d-Ser *VanN*-type transferable vancomycin resistance in *Enterococcus faecium*. Antimicrob Agents Chemother 55:4606–12.

6. Depardieu F, Perichon B, Courvalin P. 2004. Detection of the van alphabet and identification of enterococci and staphylococci at the species level by multiplex PCR. J Clin Microbiol 42:5857–60.

7. Xu X, Lin D, Yan G, Ye X, Wu S, Guo Y, Zhu D, Hu F, Zhang Y, Wang F, Jacoby GA, Wang M. 2010. *vanM*, a new glycopeptide resistance gene cluster found in *Enterococcus faecium*. Antimicrob Agents Chemother 54:4643–7.

8. Courvalin P. 2006. Vancomycin resistance in gram-positive cocci. Clin Infect Dis 42 Suppl 1:S25–34.

9. Top J, Willems R, Bonten M. 2008. Emergence of CC17 *Enterococcus faecium*: from commensal to hospital-adapted pathogen. FEMS Immunol Med Microbiol 52:297–308.

10. Ridwan B, Mascini, E., van der Reijden, N., Verhoef, J., Bonten, M. 2002. What action should be taken to prevent spread of vancomycin resistant enterococci in European hospitals? British Medical Journal 324:666–668.

11. Zirakzadeh A, Patel R. 2006. Vancomycin-resistant enterococci: colonization, infection, detection, and treatment. Mayo Clin Proc 81:529–36.

12. Stinear TP, Olden DC, Johnson PDR, Davies JK, Grayson ML. 2001. Enterococcal *vanB* resistance locus in anaerobic bacteria in human faeces. Lancet 357:855–856.

13. Howden BP, Holt KE, Lam MM, Seemann T, Ballard S, Coombs GW, Tong SY, Grayson ML, Johnson PD, Stinear TP. 2013. Genomic insights to control the emergence of vancomycin-resistant enterococci. MBio 4:e00412–3.

14. Simner PJ, Adam H, Baxter M, McCracken M, Golding G, Karlowsky JA, Nichol K, Lagace-Wiens P, Gilmour MW, Canadian Antimicrobial Resistance A, Hoban DJ, Zhanel GG. 2015. Epidemiology of vancomycin-resistant enterococci in Canadian hospitals (CANWARD study, 2007 to 2013). Antimicrob Agents Chemother 59:4315–7.

15. Werner G, Coque TM, Hammerum AM, Hope R, Hryniewicz W, Johnson A, Klare I, Kristinsson KG, Leclercq R, Lester CH, Lillie M, Novaris C, Olsson-Liljequist B, Leixe LV, Sadowy E, Simonsen GS, Top J, Vuopio-Varkila J, Willems RJ, Witte W, Woodford N. 2008. Emergence and spread of vancomycin resistance among enterococci in Europe. EuroSurveillance 13:1–11.

16. Kamarulzaman A, Tosolini, F.A., Boquest, A.L., Geddes, Richards, M.J. 1995. Vancomycin-resistant *Enterococcus faecium* in a liver transplant recipient. Aust NZ J 25:560.

17. Coombs GW, Pearson JC, Le T, Daly DA, Robinson JO, Gottlieb T, Howden BP, Johnson PD, Bennett CM, Stinear TP, Turnidge JD, on behalf of the Australian Group on Antimicrobial Resistance. 2014. Australian Enterococcal Sepsis Outcome Progamme, 2011. Commun Dis Intell Q Rep 38:E247–52.

18. Christiansen K TJ, Gottlieb T, Coombs G, Bell J, George N, Pearson J. on behalf of the Australian Group for Antimicrobial Resistance. 2010. Antimicrobial susceptibility and VRE characterisation report of enterococcus isolates from the Australian Group on Antimicrobial Resistance (AGAR): 2010 Surveillance Report.

19. Coombs GW PJ, Daley DA, Le T, Robinson O, Gottlieb T, Howden BP, Johnson PDR, Bennett CM, Stinear TP, Turnidge J. on behalf of the Australian Group on Antimicrobial Resistance. 2013. Australian Group on Antimicrobial Resistance (AGAR) Australian Enterococcal Sepsis Outcome Programme (AESOP) Annual Report 2013.

20. Coombs GW, Pearson JC, Daley DA, Le T, Robinson OJ, Gottlieb T, Howden BP, Johnson PD, Bennett CM, Stinear TP, Turnidge JD, Australian Group on Antimicrobial R. 2014. Molecular epidemiology of enterococcal bacteremia in Australia. J Clin Microbiol 52:897–905.

21. Coombs GW Daley DA, Lee YT, Pang S, Pearson JC, Robinson O, Johnson PDR, Kotsanas D, Bell JM, Turnidge J. on behalf of the Australian Group on Antimicrobial Resistance. 2014. Australian Group on Antimicrobial Resistance (AGAR) Australian Enterococcal Sepsis Outcome Programme (AESOP) Annual Report 2014.

22. Coombs GW Daley DA. on behalf of the Australian Group for Antimicrobial Resistance GoAR. 2015. Australian Enterococcal Sepsis Outcome Program (AESOP) 2015 Final Report.

23. Buultjens AH, Lam MM, Ballard S, Monk IR, Mahony AA, Grabsch EA, Grayson ML, Pang S, Coombs GW, Robinson JO, Seemann T, Johnson PD, Howden BP, Stinear TP. 2017. Evolutionary origins of the emergent ST796 clone of vancomycin resistant *Enterococcus faecium*. PeerJ 5:e2916.

24. Coombs GW, Daley D, on behalf of the Australian Group for Antimicrobial Resistance. 2017. Australian Enterococcal Sepsis Outcome Program (AESOP) 2016.

25. van Hal SJ, Espedido BA, Coombs GW, Howden BP, Korman TM, Nimmo GR, Gosbell IB, Jensen SO. 2017. Polyclonal emergence of *vanA* vancomycin-resistant *Enterococcus faecium* in Australia. J Antimicrob Chemother 72:998–1001.

26. Bolger AM, Lohse M, Usadel B. 2014. Trimmomatic: a flexible trimmer for Illumina sequence data. Bioinformatics 30:2114–20.

27. Wood D, Salzberg, SL. 2014. Kraken: ultrafast metagenomic sequence classification using exact alignments. Genome Biol 15.

28. Bankevich A, Nurk S, Antipov D, Gurevich AA, Dvorkin M, Kulikov AS, Lesin VM, Nikolenko SI, Pham S, Prjibelski AD, Pyshkin AV, Sirotkin AV, Vyahhi N, Tesler G, Alekseyev MA, Pevzner PA. 2012. SPAdes: a new genome assembly algorithm and its applications to single-cell sequencing. J Comput Biol 19:455–77.

29. Zhang Z, Schwartz S, Wagner L, Miller W. 2000. A greedy algorithm for aligning DNA sequences. J Comput Biol 7:203–14.

30. Chen L, Zheng D, Liu B, Yang J, Jin Q. 2016. VFDB 2016: hierarchical and refined dataset for big data analysis--10 years on. Nucleic Acids Res 44:D694–7.

31. Li H. 2013. Aligning sequence reads, clone sequences and assembly contigs with BWA-MEM. arXiv:1303.3997v2.

32. Garrison E, Marth, G. 2012. Haplotype-based variant detection from short-read sequencing. arXiv:1207.3907v2.

33. Robinson JT, Thorvaldsdottir H, Winckler W, Guttman M, Lander ES, Getz G, Mesirov JP. 2011. Integrative genomics viewer. Nat Biotechnol 29:24–6.

34. Didelot X, Wilson DJ. 2015. ClonalFrameML: efficient inference of recombination in whole bacterial genomes. PLoS Comput Biol 11:e1004041.

35. Page AJ, Taylor, B., Delaney, A.J., Soares, J., Seemann, T., Keane, J.A., Harris, S.R. 2016. SNP-sites: rapid efficient extraction of SNPs from multi-FASTA alignments. Microb Genom doi:10.1099/mgen.0.000056.

36. Stamatakis A. 2014. RAxML version 8: a tool for phylogenetic analysis and post-analysis of large phylogenies. Bioinformatics 30:1312–3.

37. Yu G, Smith DK, Zhu H, Guan Y, Lam TT-Y, McInerny G. 2017. ggtree: anrpackage for visualization and annotation of phylogenetic trees with their covariates and other associated data. Methods Ecol Evol 8:28–36.

38. Cheng L, Connor TR, Siren J, Aanensen DM, Corander J. 2013. Hierarchical and spatially explicit clustering of DNA sequences with BAPS software. Mol Biol Evol 30:1224–8.

39. Koren S, Walenz BP, Berlin K, Miller JR, Bergman NH, Phillippy AM. 2017. Canu: scalable and accurate long-read assembly via adaptive k-mer weighting and repeat separation. Genome Res 27:722–736.

40. Gurevich A, Saveliev V, Vyahhi N, Tesler G. 2013. QUAST: quality assessment tool for genome assemblies. Bioinformatics 29:1072–5.

41. Seemann T. 2014. Prokka: rapid prokaryotic genome annotation. Bioinformatics 30:2068–9.

42. Kearse M, Moir R, Wilson A, Stones-Havas S, Cheung M, Sturrock S, Buxton S, Cooper A, Markowitz S, Duran C, Thierer T, Ashton B, Meintjes P, Drummond A. 2012. Geneious Basic: an integrated and extendable desktop software platform for the organization and analysis of sequence data. Bioinformatics 28:1647–9.

43. Lebreton F, van Schaik W, McGuire AM, Godfrey P, Griggs A, Mazumdar V, Corander J, Cheng L, Saif S, Young S, Zeng Q, Wortman J, Birren B, Willems RJ, Earl AM, Gilmore MS. 2013. Emergence of epidemic multidrug-resistant *Enterococcus faecium* from animal and commensal strains. MBio 4:e00534–13.

44. Raven KE, Gouliouris T, Brodrick H, Coll F, Brown NM, Reynolds R, Reuter S, Torok ME, Parkhill J, Peacock SJ. 2017. Complex Routes of Nosocomial Vancomycin-resistant *Enterococcus faecium* transmission revealed by genome sequencing. Clin Infect Dis 64:886–893.

45. Raven KE, Reuter S, Reynolds R, Brodrick HJ, Russell JE, Torok ME, Parkhill J, Peacock SJ. 2016. A decade of genomic history for healthcare-associated *Enterococcus faecium* in the United Kingdom and Ireland. Genome Res 26:1388–1396.

46. Pinholt M, Gumpert H, Bayliss S, Nielsen JB, Vorobieva V, Pedersen M, Feil E, Worning P, Westh H. 2017. Genomic analysis of 495 vancomycin-resistant *Enterococcus faecium* reveals broad dissemination of a vanA plasmid in more than 19 clones from Copenhagen, Denmark. J Antimicrob Chemother 72:40–47.

47. Nguyen LT, Schmidt HA, von Haeseler A, Minh BQ. 2015. IQ-TREE: a fast and effective stochastic algorithm for estimating maximum-likelihood phylogenies. Mol Biol Evol 32:268–74.

48. European Committee on Antimicrobial Susceptibility Testing. Breakpoint tables for interpretation of MICs and zone diameters, version 7.0. 2017. http://www.eucast.org/

49. Cox CR, Coburn, P.S., Gilmore, M.S. 2005. Enterococcal cytolysin: a novel two compotent peptide system that serves as a bacterial defense against eukaryotic and prokaryotic cells. Curr Protein & Peptide Sci 6:77–84.

50. Rice LB, Carias L, Rudin S, Vael C, Goossens H, Konstabel C, Klare I, Nallapareddy SR, Huang W, Murray BE. 2003. A potential virulence gene, *hylEfm*, predominates in *Enterococcus faecium* of clinical origin. J Infect Dis 187:508–12.

51. Galli D, Lottspeich, F., Wirth, R. 1990. Sequence analysis of Enterococcus faecalis aggregation substance encoded by the sex pheromone plasmid pAD1. Mol Microbiol 4:1365–2958.

52. Su YA, Sulavik MC, He P, Makinen KK, Makinen PL, Fiedler S, Wirth R, Clewell DB. 1991. Nucleotide sequence of the gelatinase gene (*gelE*) from *Enterococcus faecalis* subsp. *liquefaciens*. Infect Immun 59:415–20.

53. Comerlato CB, de Resende MCC, Caierao J, d’Azevedo PA. 2013. Presence of virulence factors in *Enterococcus faecalis* and *Enterococcus faecium* susceptible and resistant to vancomycin. Memorias Do Instituto Oswaldo Cruz 108:590–595.

54. Nallapareddy SR, Weinstock, G.M., Murray, B.E. 2003. Clinical isolates of *Enterococcus faecium* exhibit strain-specific collagen binding mediated by *Acm*, a new member of the MSCRAMM family. Mol Microbiol 47:1733–1747.

55. Hendrickx AP, van Luit-Asbroek M, Schapendonk CM, van Wamel WJ, Braat JC, Wijnands LM, Bonten MJ, Willems RJ. 2009. *SgrA*, a nidogen-binding LPXTG surface adhesin implicated in biofilm formation, and *EcbA*, a collagen binding MSCRAMM, are two novel adhesins of hospital-acquired *Enterococcus faecium*. Infect Immun 77:5097–106.

56. Hendrickx AP, Bonten MJ, van Luit-Asbroek M, Schapendonk CM, Kragten AH, Willems RJ. 2008. Expression of two distinct types of pili by a hospital-acquired *Enterococcus faecium* isolate. Microbiology 154:3212–23.

57. Willems RJL, Homan W, Top J, van Santen-Verheuvel M, Tribe D, Manzioros X, Gaillard C, Vandenbroucke-Grauls CMJE, Mascini EM, van Kregten E, van Embden JDA, Bonten MJM. 2001. Variant *esp* gene as a marker of a distinct genetic lineage of vancomycin resistant *Enterococcus faecium* spreading in hospitals. Lancet 357:853–855.

58. Leavis H, Top J, Shankar N, Borgen K, Bonten M, van Embden J, Willems RJL. 2004. A novel putative enterococcal pathogenicity island linked to the *esp* virulence gene of *Enterococcus faecium* and associated with epidemicity. J Bacteriol 186:672–682.

59. Carter GP, Buultjens AH, Ballard SA, Baines SL, Tomita T, Strachan J, Johnson PD, Ferguson JK, Seemann T, Stinear TP, Howden BP. 2016. Emergence of endemic MLST non-typeable vancomycin-resistant *Enterococcus faecium*. J Antimicrob Chemother 71:3367–3371.

60. Christiansen K TJ, George N, Pearson J. on behalf of the Australian Group for Antimicrobial Resistance. 2005. Antimicrobial Susceptibility Report of Enterococcus Isolates from the Australian Group on Antimicrobial Resistance (AGAR): 2005 Surveillance Report.

61. Christiansen K TJ, Gottlieb, Bell J, George N, Pearson J. on behalf of the Australian Group for Antimicrobial Resistance. 2007. Enterococcus species Survey: 2007 Antimicrobial Susceptibility Report.

62. Christiansen K TJ, Gottlieb T, bell J, George N, Pearson J. on behalf of the Australian Group for Antimicrobial Resistance. 2009. Enterococcus species Survey: 2009 Antimicrobial Susceptibility Report.

63. Bell J, Turnidge J, Coombs G, O’Brien F. 1998. Emergence and epidemiology of vancomycin-resistant enterococci in Australia. Commun Dis Intell 22:249–52.

64. Worth LJ, Slavin MA, Vankerckhoven V, Goossens H, Grabsch EA, Thursky KA. 2008. Virulence determinants in vancomycin-resistant *Enterococcus faecium vanB*: clonal distribution, prevalence and significance of *esp* and *hyl* in Australian patients with haematological disorders. J Hosp Infect 68:137–44.

65. Yang J, Jiang Y, Guo L, Ye L, Ma Y, Luo Y. 2016. Prevalence of diverse clones of vancomycin-resistant *Enterococcus faecium* ST78 in a Chinese hospital. Microb Drug Resist 22:294–300.

66. Ruzon FI, de Paula SB, Kanoshiki RL, Pereira-Santos J, Kerbauy G, Kobayashi RK, Yamauchi LM, Perugini MR, Yamada-Ogatta SF. 2010. Virulence determinants in vancomycin-resistant *Enterococcus faecium vanA* isolated from different sources at University Hospital of Londrina, Parana, Brazil. J Microbiol 48:814–21.

67. Brodrick HJ, Raven KE, Harrison EM, Blane B, Reuter S, Torok ME, Parkhill J, Peacock SJ. 2016. Whole-genome sequencing reveals transmission of vancomycin-resistant *Enterococcus faecium* in a healthcare network. Genome Med 8:4.

68. Pinholt M, Larner-Svensson H, Littauer P, Moser CE, Pedersen M, Lemming LE, Ejlertsen T, Sondergaard TS, Holzknecht BJ, Justesen US, Dzajic E, Olsen SS, Nielsen JB, Worning P, Hammerum AM, Westh H, Jakobsen L. 2015. Multiple hospital outbreaks of *vanA Enterococcus faecium* in Denmark, 2012-13, investigated by WGS, MLST and PFGE. J Antimicrob Chemother 70:2474–82.

69. Lam MM, Seemann T, Bulach DM, Gladman SL, Chen H, Haring V, Moore RJ, Ballard S, Grayson ML, Johnson PD, Howden BP, Stinear TP. 2012. Comparative analysis of the first complete *Enterococcus faecium* genome. J Bacteriol 194:2334–41.

70. Weigel LM, Clewell DB, Gill SR, Clark NC, McDougal LK, Flannagan SE, Kolonay JF, Shetty J, Killgore GE, Tenover FC. 2003. Genetic analysis of a high-level vancomycin-resistant isolate of *Staphylococcus aureus*. Science 302:1569–71.

