## Supplementary Materials for "The changing landscape of VREfm in Victoria, Australia: a State-wide genomic snapshot"

The Microbiological Diagnostic Unit Public Health Laboratory, Department of Microbiology and Immunology, The University of Melbourne, at the Peter Doherty Institute for Infection and Immunity, Victoria, Australia ^a;^ Department of Microbiology and Immunology, The University of Melbourne, at the Peter Doherty Institute for Infection and Immunity, Victoria, Australia ^b^ ; Melbourne Bioinformatics, The University of Melbourne, Victoria, Australia ^c^

**Appendix**

**Appendix**

- Supplementary Methods
- Supplementary Results
  - Tables
  - Figure legends

**Supplementary Methods**

***Reference genome assembly*.** A single isolate from each major ST (17, 80, 203, 796, 1421) was sequenced using PacBio Single Molecule Real-Time (SMRT) sequencing (Pacific Biosciences, CA). *De novo* assembly was done using Canu (v.1.3, (1)), using default settings. Alignments were evaluated visually, and where fragmented, the overlap was located and joined using Geneious (v.8.1.5, Biomatters, New Zealand). Contigs were oriented to start at *dnaA* (for the chromosome) and replication-associated genes (for plasmids). Using the SMRT Analysis System v.2.3.0.140936 (Pacific Biosciences, CA), BridgeMapper was used to align long sequences reads to the assembly to assess read support for each join and regions surrounding ribosomal RNA loci. The consensus was polished using short read Illumina data extracted from the same colony. Annotation was done using Prokka (v.1.11, (2)). Raw PacBio reads and the complete assemblies for these isolates have been deposited on the National Center for Biotechnology Information’s Sequence Read Archive under BioProject PRJNA433676.

***Verifying the presence of putative virulence genes.*** With the exception of the *esp* gene, putative virulence genes were only considered present if they were at least 95% complete on assembly, without any gaps which might affect functionality.

Regarding *esp*, this gene contains several large repetitive sequences of over 250 bp in length (3), and as such, we were not able to completely assemble it *de novo* using short-read data; while many isolates had hits to the *esp* gene using the Virulence Factor DataBase (4), the assemblies of *esp* were fragmented and had incomplete coverage. To investigate this further, we reviewed the alignment of AUSMDU0004028’s Illumina reads to its own PacBio assembly. While this gene was complete in the PacBio assembly, coverage was incomplete when aligning Illumina short-reads to its own reference (this result was found when aligning reads from many other isolates to AUSMDU0004028). As shown in **Figure A4**, this was due to the presence of multi-mapping reads which spanned several regions of the gene, indicative of repeats; as these reads cannot be accurately mapped, they are removed as part of the Nullarbor pipeline. Thus, the *esp* gene cannot not be resolved using short-read data.

Due to this technical limitation, we re-examined *esp* presence/absence in our other isolates by aligning reads to an extracted, complete *esp* gene from AUSMDU0004028, and classified isolates with at least 80% *esp* gene coverage as potentially having this gene.

***Victorian strains in context.***

To assess how our strains fit in context, we downloaded all available read data from (5-8). For consistency, reads were aligned to the ST796 AUSMDU0004028 reference genome. Two isolates with only 4.48% and 20.19% coverage to the reference genome were excluded; the remainder had > 70% genome coverage to this reference. IQ-TREE (v.1.6.1, (9)) was used to generate a maximum likelihood tree, with model selection based on the lowest Bayesian Information Criteria. 20,000 ultra-fast bootstrap replicates were done to assess confidence in the resultant phylogeny. This tree used as input for ClonalFrameML (10).

Adjusting for recombination yielded only 483 core sites; given the extremely short alignment, we are concerned regarding the robustness of this analysis. While we did generate a maximum likelihood tree using these concatenated alignments, overall structure was clearly lost, with ST796 isolates, for example, were intermixed with numerous isolates from other STs and countries. Thus, we have presented the unadjusted tree in **Figure A7**.

**Supplementary Results**

***Sequencing quality.***

All 331 isolates passed quality control on WGS: the median number of reads was 2,164,564 (IQR 1,968,520-2,481,400), and the median average base quality across these reads was 32.3 (IQR 32-32.5). Mean GC content was 38.9% (SD 0.3). Kraken (v.0.10.5, (11)) was used for speciation and assessment of potential contamination, taking into account the percent coverage of reads to the *E. faecium* reference genome, the depth of coverage across this reference, and total genome size of draft assemblies. The median genome coverage to our local ST796 reference (AUSMDU00004028) was 92.2x (IQR 87.7-93.0), with a median depth of 100x (IQR 100-126). The mean genome size was 3,000,984 bp (SD 95,913).

***Short-read assembly.***

Short-reads were assembled to identify *van* genotype, MLST and putative virulence genes. The average number of contigs per assembly was 323 (SD 69), with a mean of 3,000,984 bp (SD 95,913). The average N50 was 46,625 (SD 12,877), and overall mean tRNA per assembly was 59 (SD 3).

***ST1421 subanalysis.***

Twenty-two isolates lacked the *pstS* allele (assigned ST1421 based on the remaining six alleles, **Table A3**). Reviewing the phylogeny based on SNPs called against an ST796 reference, these appeared highly clonal, and suggestive of transmission between patients. To provide higher resolution, we aligned reads from these isolates to a closer, complete ST1421 reference genome (AUSMDU00004167). Overall, there were a median of 122 core pairwise SNPs between the ST1421 isolates (IQR 12-144, range 0-274). Ten pairs of isolates were within one SNP of one another, with nine of these from the same hospital (representing seven unique patients). All were from the same healthcare network. One patient, R, was previously positive for ST80 VREfm, suggesting either multiple infection at baseline or reinfection with this strain.

***Within-host diversity.***

There is limited data on within-host diversity of VRE; when examining repeat isolates from three VRE-positive nursing home residents, Brodrick *et al.* (12) found up to 6 SNPs between isolates over a six month period. To examine within-host diversity in our dataset, and also assess for potential re-infection, we examined the number of SNPs between repeat isolates from the same patients.

Twenty-seven patients had >1 sample positive for *E. faecium* (range 2-5 per patient). All repeat samples were collected within the same hospital as the original sample, except for one where isolates were collected at two different hospitals within the same healthcare network (**Table A2**). Twenty-two patients had repeat samples with isolates that had the same ST, with a median of 0 core SNPs, and a maximum of 40 SNPs between any two isolates (**Figure A1**). Two patients (E and O, **Table A2**) had isolates that accounted for most of the diversity. Patient E had three isolates collected over ~one week. Two were clinical samples, with isolates that were only one SNP apart from one another, while the third isolate was collected from a screening swab from a different anatomical location and was 14-15 SNPs from clinical isolates with a slightly different resistome. Patient O likewise had three positive screening samples collected over a total of 2 weeks. The earliest isolate was 40 SNPs from others; the remaining two isolates, collected a week apart, had identical core genomes to one another.

Six patients had repeat samples with isolates of different STs (**Table A2**), indicating either re-infection with a novel ST or mixed infection at baseline. As expected, these isolates had much higher diversity, with 171-334 core SNPs between any two isolates (**Figure A1**).

All three repeat pairs from ST1421 were from a single patient, and had identical core genomes.

One patient (patient A) had a VSEfm isolate and a *vanA-*positive isolate collected from two blood samples in the same day (**Table A2**). These isolates differed by only four core SNPs, suggesting that the latter isolate had lost the *vanA* plasmid during passage. We have observed this infrequently in other VREfm projects at the Microbiological Diagnostic Unit Public Health Laboratory (unpublished data).

***Other antibiotic resistance genes on the vanA-harbouring plasmid.***

The *ermB* gene, conferring resistance to erythromycin, was identified on all *vanA-*harbouring plasmids. *ant(6)-a1,* which confers resistance to aminoglycosides, was present on the *vanA-*harbouring plasmids from ST 78, ST203, and ST796 isolates, while ­*aph(3’)-III* was present on the *vanA*-harbouring plasmids from ST78, 796, and ST1421, respectively. *cat*pC221, a chloramphenicol acetyltransferase, was also present on the plasmid from the ST203 isolate (**Figure A5**).

**Supplementary Tables**

**Table A1.** **Single molecule, real-time sequencing assembly quality**

| Isolate | ST | Total size, bp^a^ | Total number of contigs^a^ | N50, bp^a^ | GC content (%)^a^ | Number of tRNAs^b^ | Contig | Contig size, bp^c^ |
| --- | --- | --- | --- | --- | --- | --- | --- | --- |
| AUSMDU00004024 | 203 | 3,170,680 | 3 | 2,863,087 | 37.77 | 70 | Chromosome | 2,863,087 |
|  |  |  |  |  |  |  | Plasmid | 259,384 |
|  |  |  |  |  |  |  | Plasmid | 49,013^d^ |
| AUSMDU00004028 | 796 | 3,185,609 | 5 | 2,855,729 | 37.85 | 72 | Chromosome | 2,855,729 |
|  |  |  |  |  |  |  | Plasmid | 184,337 |
|  |  |  |  |  |  |  | Plasmid | 54,737 |
|  |  |  |  |  |  |  | Plasmid | 53,654 |
|  |  |  |  |  |  |  | Plasmid | 37,772 ^d^ |
| AUSMDU00004142 | 80 | 3,202,267 | 5 | 2,912,017 | 37.77 | 69 | Chromosome | 2,912,017 |
|  |  |  |  |  |  |  | Plasmid | 168,778 |
|  |  |  |  |  |  |  | Plasmid | 54,738 |
|  |  |  |  |  |  |  | Plasmid | 36,024 |
|  |  |  |  |  |  |  | Plasmid | 31,222 ^d^ |
| AUSMDU00004167 | 1421 | 3,200,573 | 4 | 2,883,877 | 37.72 | 69 | Chromosome | 2,883,877 |
|  |  |  |  |  |  |  | Plasmid | 207,001 |
|  |  |  |  |  |  |  | Plasmid | 63,221 |
|  |  |  |  |  |  |  | Plasmid | 47,249 ^d^ |
| AUSMDU00004055 | 78 | 3,177,207 | 6 | 2,731,844 | 37.65 | 68 | Chromosome | 2,731,844 |
|  |  |  |  |  |  |  | Plasmid | 231,024 |
|  |  |  |  |  |  |  | Plasmid | 91,057 |
|  |  |  |  |  |  |  | Plasmid | 53,613 |
|  |  |  |  |  |  |  | Plasmid | 36,341 |
|  |  |  |  |  |  |  | Plasmid | 33,328 ^d^ |

^a^QUAST (v.4.5, (13) was used to calculate these values. ^b^Prokka (https://github.com/tseemann/prokka/) was used to annotate tRNA genes. ^c^Chromosome and plasmid size were calculated from the annotated genbank files. Totals of chromosome + plasmid may not sum to exactly that listed in column 3, due to differences in the tools used. ^d^Indicates the plasmid with the *vanA* gene.

**Table A2.** **Characteristics of repeat samples from the same patients**

| Patient | Isolate | Date collected | Hospital network | Hospital | Blood-stream infection | MLST | *van* gene |
| --- | --- | --- | --- | --- | --- | --- | --- |
| **A** | AUSMDU00004142 | 30/11/2015 | H4 | HCN4 | Yes | **80** | ***vanA*** |
|  | AUSMDU00004137 | 30/11/2015 | H4 | HCN4 | Yes | **80** | **- ^a^** |
| B | AUSMDU00004271 | 18/11/2015 | H2 | HCN3 | No | 796 | *vanB* |
|  | AUSMDU00004272 | 20/11/2015 | H2 | HCN3 | No | 796 | *vanB* |
| C | AUSMDU00004283 | 30/11/2015 | H2 | HCN3 | No | 796 | *vanB* |
|  | AUSMDU00004287 | 4/12/2015 | H2 | HCN3 | No | 796 | *vanB* |
|  | AUSMDU00004295 | 7/12/2015 | H2 | HCN3 | Yes | 796 | *vanB* |
| D | AUSMDU00004288 | 5/12/2015 | H2 | HCN3 | No | 796 | *vanB* |
|  | AUSMDU00004297 | 7/12/2015 | H2 | HCN3 | No | 796 | *vanB* |
| E | AUSMDU00004277 | 27/11/2015 | H2 | HCN3 | No | 203 | *vanB* |
|  | AUSMDU00004282 | 30/11/2015 | H2 | HCN3 | No | 203 | *vanB* |
|  | AUSMDU00004286 | 2/12/2015 | H2 | HCN3 | No | 203 | *vanB* |
| **F** | AUSMDU00004284 | 2/12/2015 | H2 | HCN3 | No | **203** | ***vanA*** |
|  | AUSMDU00004300 | 9/12/2015 | H2 | HCN3 | No | **796** | ***vanB*** |
| G | AUSMDU00004154 | 8/12/2015 | H11 | HCN8 | No | 796 | *vanB* |
|  | AUSMDU00004256 | 8/12/2015 | H11 | HCN8 | No | 796 | *vanB* |
| H | AUSMDU00004012 | 13/11/2015 | H16 | HCN15 | No | 796 | *vanB* |
|  | AUSMDU00004010 | 13/11/2015 | H16 | HCN15 | No | 796 | *vanB* |
| I | AUSMDU00004091 | 20/11/2015 | H18 | HCN16 | No | 203 | *vanA* |
|  | AUSMDU00004125 | 30/11/2015 | H18 | HCN16 | No | 203 | *vanA* |
| **J** | AUSMDU00004127 | 28/11/2015 | **H25** | HCN17 | No | **80** | ***vanA*** |
|  | AUSMDU00004254 | 7/12/2015 | **H40** | HCN17 | No | **796** | ***vanB*** |
| K | AUSMDU00004037 | 22/11/2015 | H17 | HCN14 | Yes | 796 | *vanB* |
|  | AUSMDU00004066 | 23/11/2015 | H17 | HCN14 | No | 796 | *vanB* |
| L | AUSMDU00004075 | 27/11/2015 | H3 | HCN2 | No | 80 | *vanA* |
|  | AUSMDU00004088 | 28/11/2015 | H3 | HCN2 | No | 80 | *vanA* |
| M | AUSMDU00004079 | 25/11/2015 | H8 | HCN2 | No | 796 | *vanB* |
|  | AUSMDU00004102 | 30/11/2015 | H8 | HCN2 | No | 796 | *vanB* |
| N | AUSMDU00004081 | 22/11/2015 | H8 | HCN2 | No | 796 | *vanB* |
|  | AUSMDU00004087 | 27/11/2015 | H8 | HCN2 | Yes | 796 | *vanB* |
| O | AUSMDU00004201 | 23/11/2015 | H1 | HCN1 | No | 17 | *vanB* |
|  | AUSMDU00004214 | 1/12/2015 | H1 | HCN1 | No | 17 | *vanB* |
|  | AUSMDU00004231 | 8/12/2015 | H1 | HCN1 | No | 17 | *vanB* |
| **P** | AUSMDU00004180 | 13/11/2015 | H1 | HCN1 | No | **1429** | *vanB* |
|  | AUSMDU00004197 | 20/11/2015 | H1 | HCN1 | No | **1429** | *vanB* |
|  | AUSMDU00004203 | 24/11/2015 | H1 | HCN1 | No | **1429** | *vanB* |
|  | AUSMDU00004216 | 1/12/2015 | H1 | HCN1 | No | **1429** | *vanB* |
|  | AUSMDU00004233 | 8/12/2015 | H1 | HCN1 | No | **17** | *vanB* |
| **Q** | AUSMDU00004165 | 17/11/2015 | H1 | HCN1 | No | **796** | *vanB* |
|  | AUSMDU00004212 | 30/11/2015 | H1 | HCN1 | No | **117** | *vanB* |
| **R** | AUSMDU00004172 | 10/11/2015 | H1 | HCN1 | No | **80** | ***vanB*** |
|  | AUSMDU00004199 | 20/11/2015 | H1 | HCN1 | No | **80** | ***vanB*** |
|  | AUSMDU00004206 | 24/11/2015 | H1 | HCN1 | No | **1421** | ***vanA*** |
| S | AUSMDU00004173 | 10/11/2015 | H1 | HCN1 | No | 796 | *vanB* |
|  | AUSMDU00004177 | 13/11/2015 | H1 | HCN1 | No | 796 | *vanB* |
| T | AUSMDU00004190 | 17/11/2015 | H1 | HCN1 | No | 796 | *vanB* |
|  | AUSMDU00004196 | 20/11/2015 | H1 | HCN1 | No | 796 | *vanB* |
| U | AUSMDU00004218 | 3/12/2015 | H1 | HCN1 | No | 192 | *vanB* |
|  | AUSMDU00004229 | 7/12/2015 | H1 | HCN1 | No | 192 | *vanB* |
| V | AUSMDU00004166 | 29/11/2015 | H1 | HCN1 | No | 1421 | *vanA* |
|  | AUSMDU00004167 | 30/11/2015 | H1 | HCN1 | No | 1421 | *vanA* |
|  | AUSMDU00004219 | 3/12/2015 | H1 | HCN1 | No | 1421 | *vanA* |
| W | AUSMDU00004186 | 15/11/2015 | H1 | HCN1 | No | **?new** | *vanB* |
|  | AUSMDU00004223 | 6/12/2015 | H1 | HCN1 | No | **796** | *vanB* |
| X | AUSMDU00004189 | 17/11/2015 | H1 | HCN1 | No | 796 | *vanB* |
|  | AUSMDU00004195 | 20/11/2015 | H1 | HCN1 | No | 796 | *vanB* |
|  | AUSMDU00004202 | 24/11/2015 | H1 | HCN1 | No | 796 | *vanB* |
| Y | AUSMDU00004181 | 13/11/2015 | H1 | HCN1 | No | 796 | *vanB* |
|  | AUSMDU00004198 | 20/11/2015 | H1 | HCN1 | No | 796 | *vanB* |
| BB | AUSMDU00004315 | 10/11/2015 | H6 | HCN6 | No | 796 | *vanB* |
|  | AUSMDU00004320 | 20/11/2015 | H6 | HCN6 | No | 796 | *vanB* |
| CC | AUSMDU00004246 | 23/11/2015 | H13 | HCN7 | No | 796 | *vanB* |
|  | AUSMDU00004247 | 26/11/2015 | H13 | HCN7 | No | 796 | *vanB* |

**^a^**This isolate is only 4 core SNPs from the other, *vanA*-positive isolate from the same patient. As both were also blood-stream samples, this suggests the *vanA*-carrying plasmid may have been initially present, but lost during culture.

**Table A3. Multi-locus sequence typing alleles for all 22 *pstS*(-) isolates in this dataset, corresponding to ST1421**

Alleles

*atpA*(1)

*ddl*(1)

*gdh*(1)

*purK*(1)

*gyd*(1)

*pstS*(-)

*adk*(1)

**Table A4.** **Closest nucleotide match to *vanA*-harbouring plasmids from VRE in Victoria using BLASTN (v.2.8.0+, (14))**

| Isolate corresponding to the vanA-harbouring plasmid | ST | Plasmid size, bp ^a^ | Top BLASTN hit | Query cover (%) | Identity (%) | NCBI Accession | Submitter | Citation |
| --- | --- | --- | --- | --- | --- | --- | --- | --- |
| AUSMDU00004055 | 78 | 33,328 | Efm V24 plasmid pHvH-V24 | 100 | 99 | KX574671.1 | Hvidovre Hospital, Denmark | (8) |
| AUSMDU00004028 | 796 | 37,772 | Efm V24 plasmid pHvH-V24 | 91 | 99 | KX574671.1 | Hvidovre Hospital, Denmark | (8) |
| AUSMDU00004142 | 80 | 31,222 | Efm V24 plasmid pHvH-V24 | 90 | 99 | KX574671.1 | Hvidovre Hospital, Denmark | (8) |
| AUSMDU00004024 | 203 | 49,013 | Efm0038 plasmid pJEG043 | 60 | 99 | KX810026.1 | Western Sydney University, Australia | (15) |
| AUSMDU00004167 | 1421 | 47,249 | Ef_DMG1500501 ^b^ plasmid 4 | 100 | 99 | LT603681.1 | University of Melbourne, Australia | (16) |

^a^ Plasmid size was calculated from the annotated Genbank files. ^b^ Communication with genome authors indicates this isolate is actually ‘Ef_DMG1500801’ as described in the cited manuscript.

**Supplementary Figure Legends.**

**Figure A1. Within-host diversity of *E. faecium***

Each black dot represents a pairwise comparison within a single patient. The median of all pairwise comparisons by ST is shown in red. Except one, all isolates from the same patients had a *van* gene.

**Figure A2. Recombination blocks identified in Victorian *E. faecium***

6,497 recombination-adjusted core SNPs from all 331 isolates (321 VREfm and 10 VSEfm) were concatenated, and used to produce a maximum likelihood tree in RAxML (17) under a General Time Reversible model with gamma rate heterogeneity. One thousand bootstrap replicates were performed to assess confidence in the phylogeny. To allow for visualization, the branch with AUSMDU00004157 (ST54), which was >3,800 SNPs from all other isolates, has been truncated in the above. Branches with bootstrap support >90 are shown. As many branches had very low bootstrap support, hierarchical Bayesian Analysis of Population Structure was used to further validate the clusters identified. Main BAPS groups are shown. The reference is identified with an arrow. Recombination blocks are highlighted in grey; their precise position according to the reference genome is indicated on the X-axis.

**Figure A3. Distribution of Victorian *E. faecium* across healthcare networks**

6,497 recombination-adjusted core SNPs from all 331 isolates (321 VREfm and 10 VSEfm) were concatenated, and used to produce a maximum likelihood tree in RAxML (17) under a General Time Reversible model with gamma rate heterogeneity. One thousand bootstrap replicates were performed to assess confidence in the phylogeny. Branches with bootstrap support >90 are shown. As many branches had very low bootstrap support, hierarchical Bayesian Analysis of Population Structure was used to further validate the clusters identified. Main BAPS groups are shown. The reference is identified with an arrow. HCN = healthcare network.  ‘.’ indicates missing data on HCN, while N/A indicates the isolate was collected at a doctor’s or dental office.

**Figure A4.** **Coverage of Illumina reads from AUSMDU0000428 to the *esp* gene locus, using the same isolate’s complete Single Molecule, Real-Time sequencing assembly as reference**

Illumina NextSeq 150bp paired-end reads from AUSMDU0000428 were aligned to the polished, corrected and annotated complete assembly of the same genome, and visualized using Integrative Genomics Viewer (v.2.3.88, (18)). The alignment has been zoomed in on the coordinates of the *esp* gene. These figures are representative of alignments to *esp* obtained for other isolates in this dataset.

**A.** Trimmed reads were aligned using default settings with BWA MEM (v.0.7.16a-r1181, (19)). Multi-mapping reads, i.e., reads that align to >1 locus, are indicated in white.

**B.** The same alignment is shown after running snippy (<https://github.com/tseemann/snippy/blob/master/bin/snippy)>, which is part of the Nullarbor pipeline; snippy, by default, excludes multi-mapping reads because from the alignment because these cannot be accurately mapped, thus giving the appearance of zero depth of coverage over these repetitive regions.

**Figure A5. v*anA*-harbouring plasmids from isolates representing each sequence type identified in Victoria**

PacBio Single Molecule Real-Time sequencing was performed on a single *vanA-*positive isolate from each of the main STs identified (ST 78, 80, 203, 796, 1421). Plasmids harbouring *vanA* were initially identified using ABRicate (https://github.com/tseemann/abricate), and subsequently annotated using the Enterococcus database (https://github.com/tseemann/prokka/blob/master/db/genus/Enterococcus). Plasmids were then circularized and visualized in Geneious (v.9.1.7, (20)). Open reading frames (ORFs) were also predicted in Geneious, with a minimum of 100 bp required. Annotated genes are shown in red on the outmost circle, and predicted ORFs are orange. GC content is shown in blue, while AT content is shown in green. * represents a hypothetical protein.

**Figure A6. Homology with closely-related published *vanA*-harbouring plasmids identified in Table A4**

PacBio Single Molecule Real-Time sequencing was performed on a total of five *vanA-*positive isolates, each representing one of the predominant ST types identified (ST 78, 80, 203, 796 and 1421). The *vanA*-harbouring plasmids were identified as described, with annotation done using prokka (v.1.12) based on the Enterococcus database (<https://github.com/tseemann/prokka/blob/master/db/genus/Enterococcus)>. BLASTn (14) was used to identify published plasmids with the greatest sequence homology (**Table A4)**. Published plasmids (from (8, 15, 16)) were likewise annotated with prokka, and alignments were generated using progressiveMauve (build date 2014-12-19). The *vanA* is indicated on each alignment using the black marker.

**Figure A7. Victorian *E. faecium* in context with published isolates**

A phylogenetic tree of the 331 Australian isolates, plus 877 isolates from the literature (5-8), based on a concatenated SNP alignment with 111,858 positions. IQ-TREE (v.1.6.1, (9)) was used to generate a maximum likelihood tree, with 20,000 ultra-fast bootstrap replicates. The tree is rooted at the midpoint. For clarity, European countries with fewer than 10 isolates are shown as ‘Other, Europe’. The ‘United Kingdom’ includes isolates that were listed only as United Kingdom, as well as isolates from Wales, Scotland and England. As isolates from the Republic of Ireland and Northern Ireland were not specifically distinguished from one another, these are simply coloured as ‘Ireland’ above. Where only one isolate was available from a continent, these were coloured by the continent. Isolates were classified as ‘Clinical’, ‘Non-clinical’, ‘Animal’, etc., per previous publications. If an isolate had no previous annotation, but was taken from human blood, it was also classified as ‘Clinical’ for this analysis. This tree was visualizing using FigTree (v.1.4.2). Branch lengths have been transformed proportionally for easier visualization of tree structure. The most represented ST among *vanA* and *vanB*-VREfm is indicated.

**References**

1.	Koren S, Walenz BP, Berlin K, Miller JR, Bergman NH, Phillippy AM. 2017. Canu: scalable and accurate long-read assembly via adaptive k-mer weighting and repeat separation. Genome Res 27:722-736.

2.	Seemann T. 2014. Prokka: rapid prokaryotic genome annotation. Bioinformatics 30:2068-9.

3.	Leavis H, Top J, Shankar N, Borgen K, Bonten M, van Embden J, Willems RJL. 2004. A novel putative enterococcal pathogenicity island linked to the *esp* virulence gene of *Enterococcus faecium* and associated with epidemicity. J Bacteriol 186:672-682.

4.	Chen L, Zheng D, Liu B, Yang J, Jin Q. 2016. VFDB 2016: hierarchical and refined dataset for big data analysis--10 years on. Nucleic Acids Res 44:D694-7.

5.	Lebreton F, van Schaik W, McGuire AM, Godfrey P, Griggs A, Mazumdar V, Corander J, Cheng L, Saif S, Young S, Zeng Q, Wortman J, Birren B, Willems RJ, Earl AM, Gilmore MS. 2013. Emergence of epidemic multidrug-resistant *Enterococcus faecium* from animal and commensal strains. MBio 4:e00412-3.

6.	Raven KE, Gouliouris T, Brodrick H, Coll F, Brown NM, Reynolds R, Reuter S, Torok ME, Parkhill J, Peacock SJ. 2017. Complex Routes of nosocomial vancomycin-resistant *Enterococcus faecium* transmission revealed by genome sequencing. Clin Infect Dis 64:886-893.

7.	Raven KE, Reuter S, Reynolds R, Brodrick HJ, Russell JE, Torok ME, Parkhill J, Peacock SJ. 2016. A decade of genomic history for healthcare-associated *Enterococcus faecium* in the United Kingdom and Ireland. Genome Res 26:1388-1396.

8.	Pinholt M, Gumpert H, Bayliss S, Nielsen JB, Vorobieva V, Pedersen M, Feil E, Worning P, Westh H. 2017. Genomic analysis of 495 vancomycin-resistant *Enterococcus faecium* reveals broad dissemination of a *vanA* plasmid in more than 19 clones from Copenhagen, Denmark. J Antimicrob Chemother 72:40-47.

9.	Nguyen LT, Schmidt HA, von Haeseler A, Minh BQ. 2015. IQ-TREE: a fast and effective stochastic algorithm for estimating maximum-likelihood phylogenies. Mol Biol Evol 32:268-74.

10.	Didelot X, Wilson DJ. 2015. ClonalFrameML: efficient inference of recombination in whole bacterial genomes. PLoS Comput Biol 11:e1004041.

11.	Wood D, Salzberg, SL. 2014. Kraken: ultrafast metagenomic sequence classification using exact alignments. Genome Biol 15.

12.	Brodrick HJ, Raven KE, Harrison EM, Blane B, Reuter S, Torok ME, Parkhill J, Peacock SJ. 2016. Whole-genome sequencing reveals transmission of vancomycin-resistant *Enterococcus faecium* in a healthcare network. Genome Med 8:4.

13.	Gurevich A, Saveliev V, Vyahhi N, Tesler G. 2013. QUAST: quality assessment tool for genome assemblies. Bioinformatics 29:1072-5.

14.	Zhang Z, Schwartz S, Wagner L, Miller W. 2000. A greedy algorithm for aligning DNA sequences. J Comput Biol 7:203-14.

15.	van Hal SJ, Espedido BA, Coombs GW, Howden BP, Korman TM, Nimmo GR, Gosbell IB, Jensen SO. 2017. Polyclonal emergence of *vanA* vancomycin-resistant *Enterococcus faecium* in Australia. J Antimicrob Chemother 72:998-1001.

16.	Carter GP, Buultjens AH, Ballard SA, Baines SL, Tomita T, Strachan J, Johnson PD, Ferguson JK, Seemann T, Stinear TP, Howden BP. 2016. Emergence of endemic MLST non-typeable vancomycin-resistant *Enterococcus faecium*. J Antimicrob Chemother 71:3367-3371.

17.	Stamatakis A. 2014. RAxML version 8: a tool for phylogenetic analysis and post-analysis of large phylogenies. Bioinformatics 30:1312-3.

18.	Robinson JT, Thorvaldsdottir H, Winckler W, Guttman M, Lander ES, Getz G, Mesirov JP. 2011. Integrative genomics viewer. Nat Biotechnol 29:24-6.

19.	Bolger AM, Lohse M, Usadel B. 2014. Trimmomatic: a flexible trimmer for Illumina sequence data. Bioinformatics 30:2114-20.

20.	Kearse M, Moir R, Wilson A, Stones-Havas S, Cheung M, Sturrock S, Buxton S, Cooper A, Markowitz S, Duran C, Thierer T, Ashton B, Meintjes P, Drummond A. 2012. Geneious Basic: an integrated and extendable desktop software platform for the organization and analysis of sequence data. Bioinformatics 28:1647-9.
