## Supplementary Materials for "The changing landscape of VREfm in Victoria, Australia: a State-wide genomic snapshot"

**Figure A1.** Within-host diversity of *E. faecium*

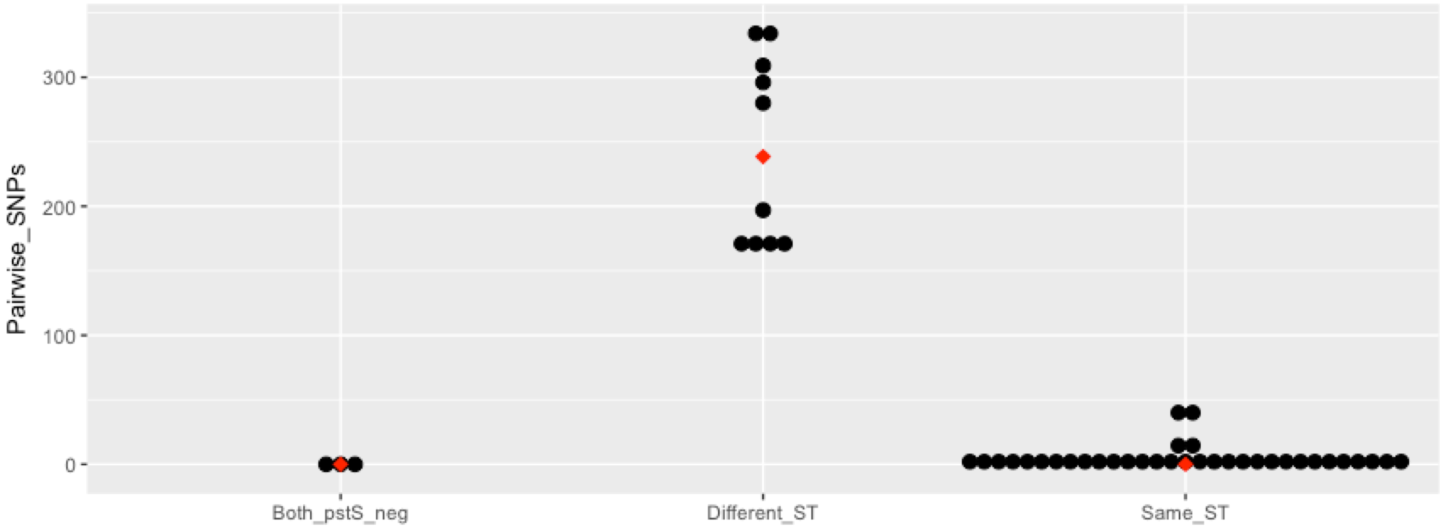


**Legend.** Each black dot represents a pairwise comparison within a single patient. The median pairwise comparisons by ST group is shown in red. Except one, all isolates from the same patients had a *van* gene.

**Figure A2.** Recombination blocks identified in Victorian *E. faecium*

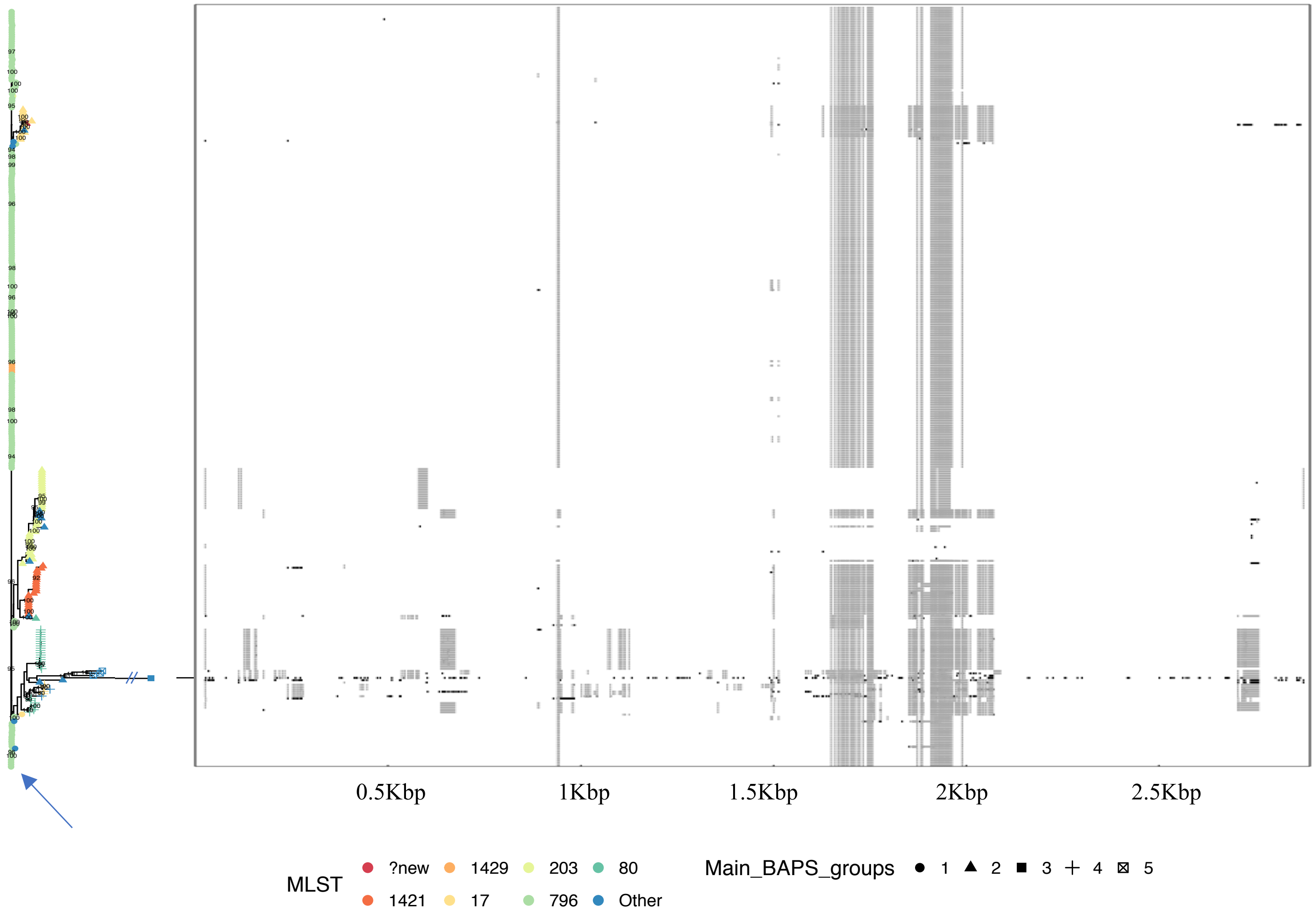


**Legend.** 6,497 recombination-adjusted core SNPs from all 331 isolates (321 VREfm and 10 VSEfm) were concatenated, and used to produce a maximum likelihood tree in RAxML (17) under a General Time Reversible model with gamma rate heterogeneity. One thousand bootstrap replicates were performed to assess confidence in the phylogeny. To allow for visualization, the branch with AUSMDU00004157 (ST54), which was >3,800 SNPs from all other isolates, has been truncated in the above. Branches with bootstrap support >90 are shown. As many branches had very low bootstrap support, hierarchical Bayesian Analysis of Population Structure was used to further validate the clusters identified. Main BAPS groups are shown. The reference is identified with an arrow. Recombination blocks are highlighted in grey; their precise position according to the reference genome is indicated on the X-axis.

**Figure A3.** Distribution of Victorian *E. faecium* across healthcare networks

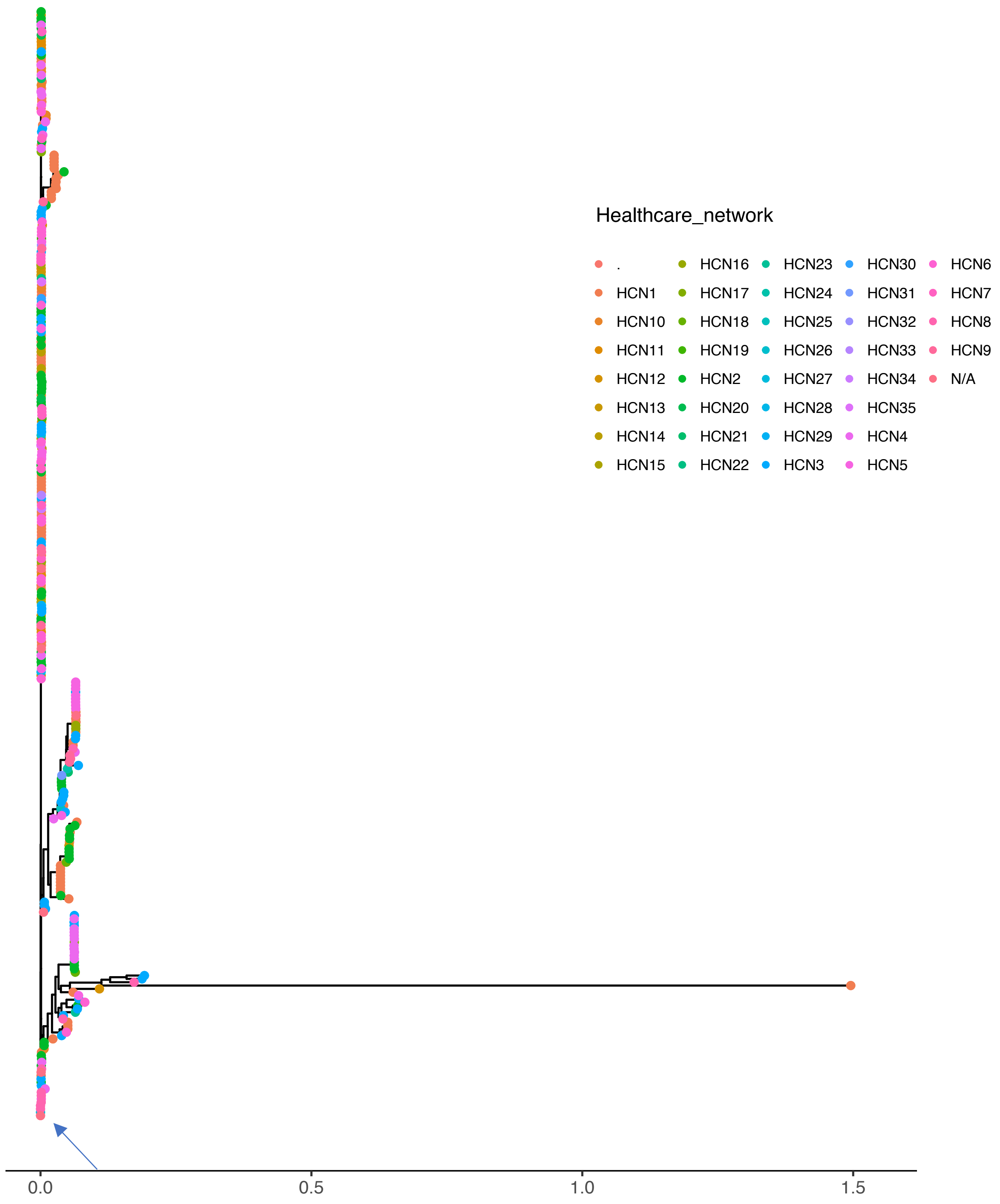


**Legend.** 6,497 recombination-adjusted core SNPs from all 331 isolates (321 VREfm and 10 VSEfm) were concatenated, and used to produce a maximum likelihood tree in RAxML (17) under a General Time Reversible model with gamma rate heterogeneity. One thousand bootstrap replicates were performed to assess confidence in the phylogeny. Branches with bootstrap support >90 are shown. As many branches had very low bootstrap support, hierarchical Bayesian Analysis of Population Structure was used to further validate the clusters identified. Main BAPS groups are shown. The reference is identified with an arrow. HCN = healthcare network. '.' indicates missing data on HCN, while N/A indicates the isolate was collected at a doctor's or dental office.

**Figure A4.** Coverage of Illumina reads from AUSMDU0000428 to the *esp* gene locus, using the same isolate's complete Single Molecule, Real-Time sequencing assembly as reference

A.

B.

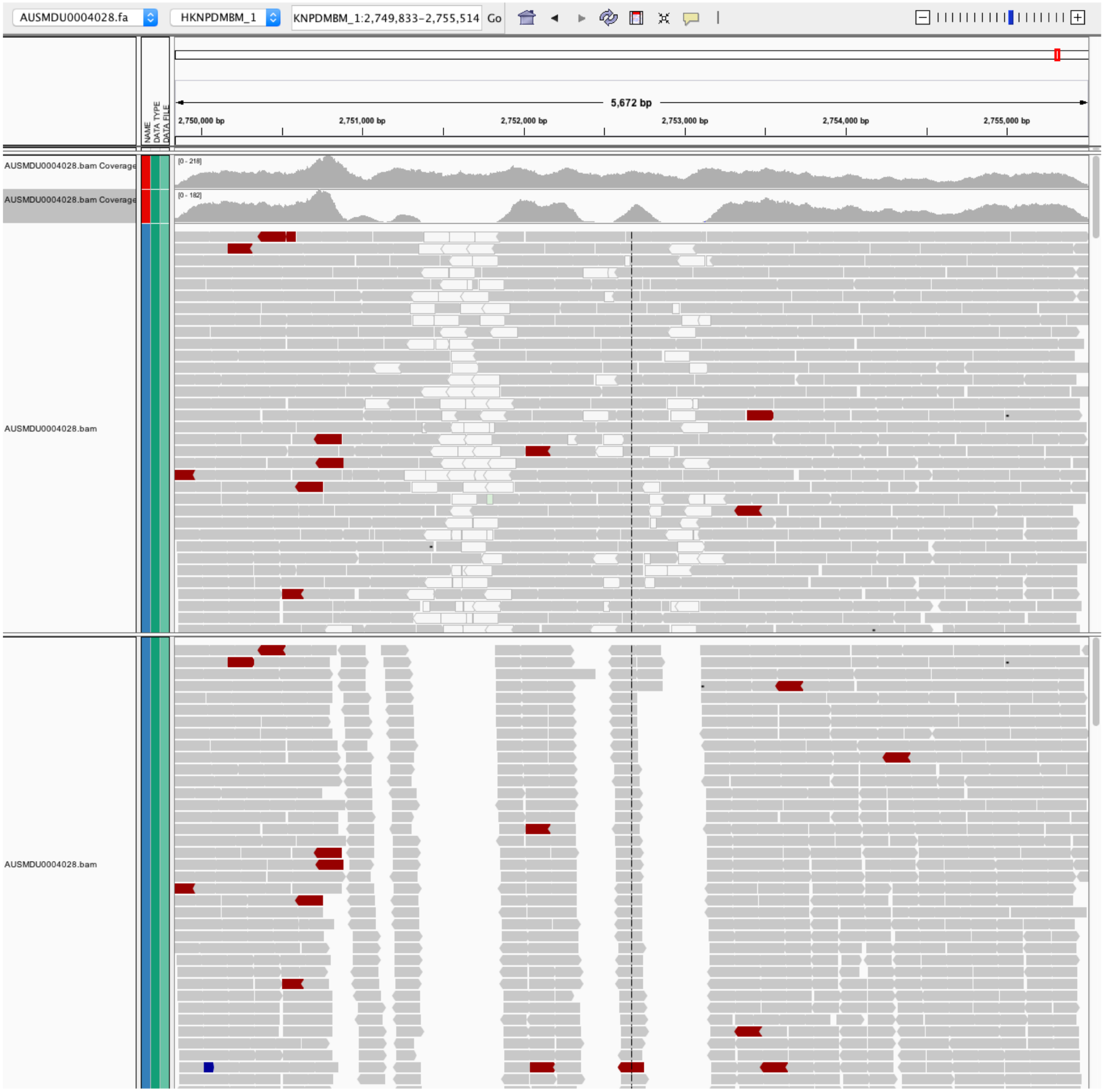


**Legend.** Illumina NextSeq 150bp paired-end reads from AUSMDU0000428 were aligned to the polished, corrected and annotated complete assembly of the same genome, and visualized using Integrative Genomics Viewer (v.2.3.88, (18)). The alignment has been zoomed in on the coordinates of the *esp* gene. These figures are representative of alignments to *esp* obtained for other isolates in this dataset.

**A.** Trimmed reads were aligned using default settings with BWA MEM (v.0.7.16a-r1181, (19)). Multi-mapping reads, i.e., reads that align to >1 locus, are indicated in white.

**B.** The same alignment is shown after running snippy (https://github.com/tseemann/snippy/blob/master/bin/snippy), which is part of the Nullarbor pipeline; snippy, by default, excludes multi-mapping reads because from the alignment because these cannot be accurately mapped, thus giving the appearance of zero depth of coverage over these repetitive regions.

**Figure A5.** *vanA*-harbouring plasmids from isolates representing each sequence type identified in Victoria

ST78

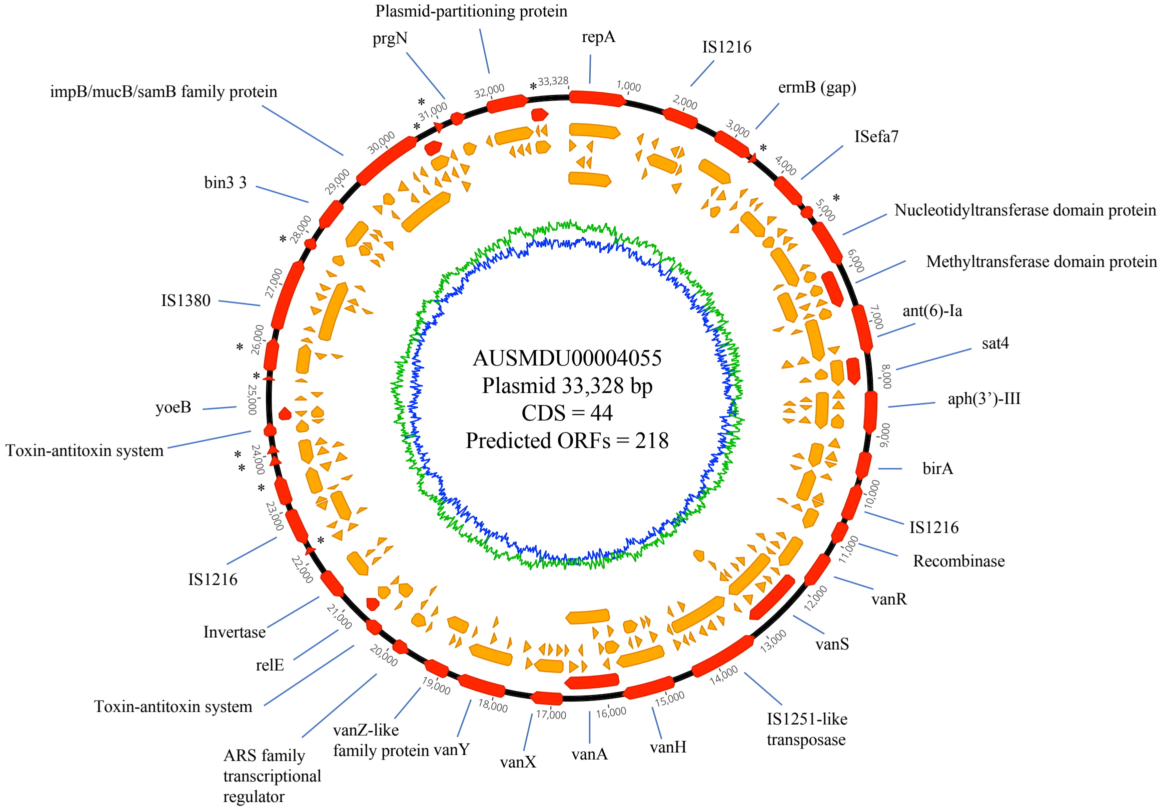


ST796

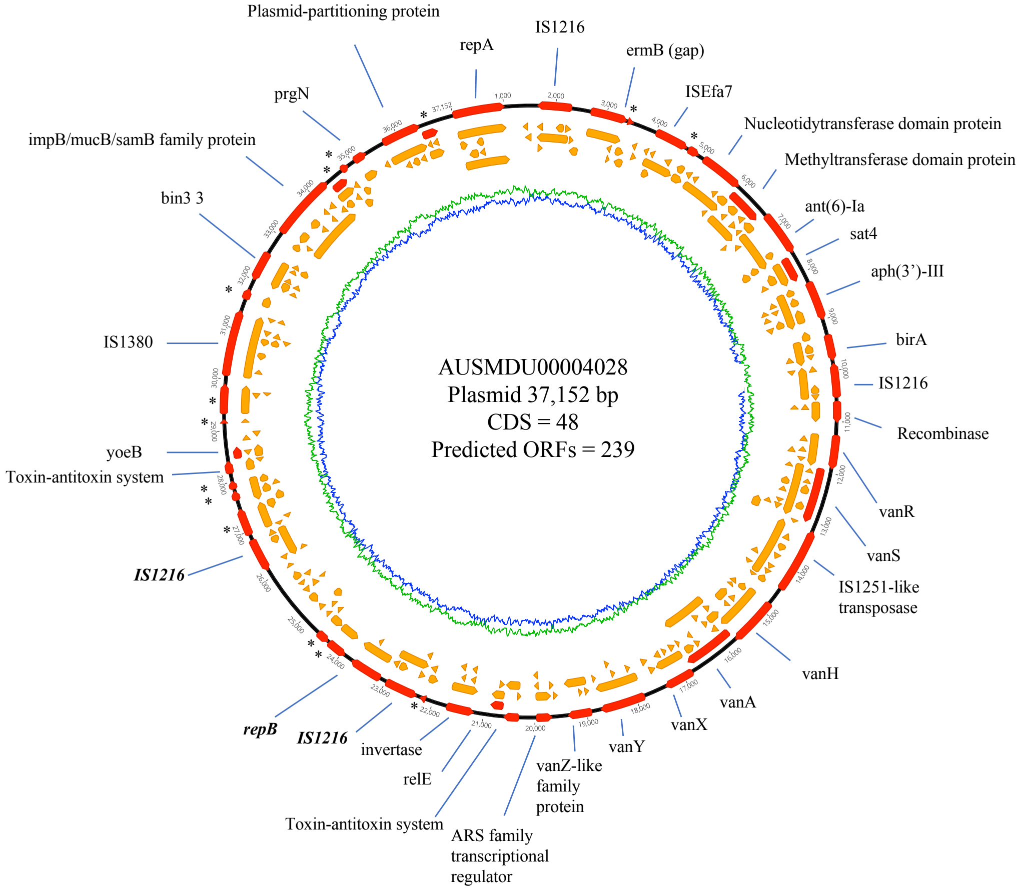

ST80

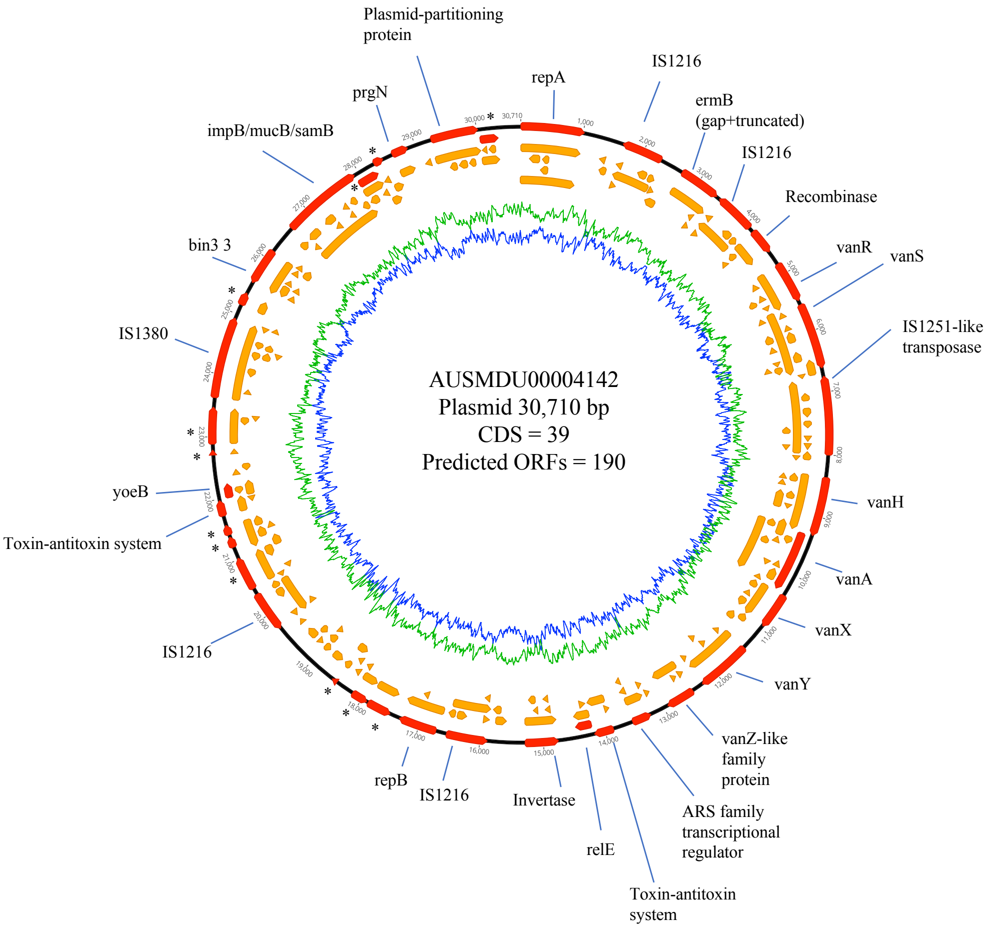
AUSMDU00004142
Plasmid 30,710 bp
CDS = 39
Predicted ORFs = 190
Plasmid-partitioning protein
repA
IS1216
ermB (gap+truncated)
IS1216
Recombinase
vanR
vanS
IS1251-like transposase
vanH
vanA
vanX
vanY
vanZ-like family protein
ARS family transcriptional regulator
Toxin-antitoxin system
relE
Invertase
IS1216
repB
IS1216
Toxin-antitoxin system
yoeB
IS1380
bin3 3
impB/mucB/samB
prgN

ST203

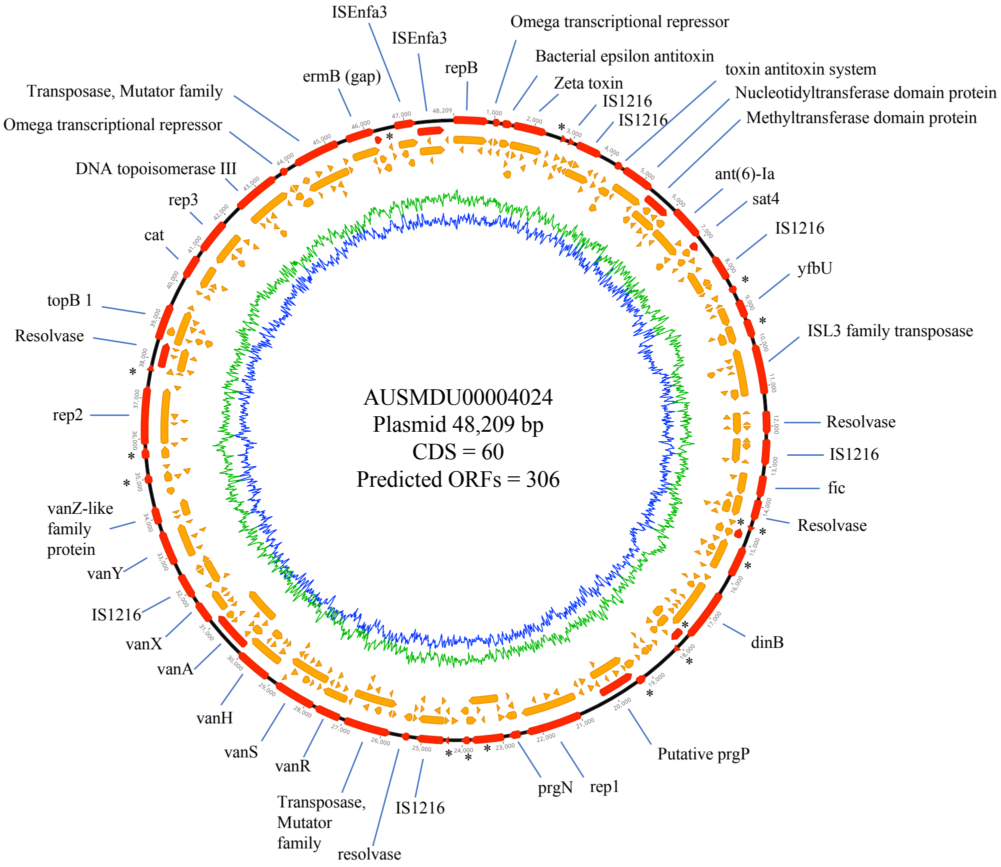
AUSMDU00004024
Plasmid 48,209 bp
CDS = 60
Predicted ORFs = 306
ISEnfa3
ISEnfa3
repB
Omega transcriptional repressor
Bacterial epsilon antitoxin
Zeta toxin
IS1216
IS1216
toxin antitoxin system
Nucleotidyltransferase domain protein
Methyltransferase domain protein
ant(6)-Ia
sat4
IS1216
yfbU
ISL3 family transposase
Resolvase
IS1216
fic
Resolvase
dinB
Putative prgP
rep1
prgN
IS1216
resolvase
Transposase, Mutator family
vanR
vanS
vanH
vanA
vanX
IS1216
vanY
vanZ-like family protein
rep2
Resolvase
topB 1
cat
rep3
DNA topoisomerase III
Omega transcriptional repressor
Transposase, Mutator family
ermB (gap)

ST1421 (*pstS*(-))

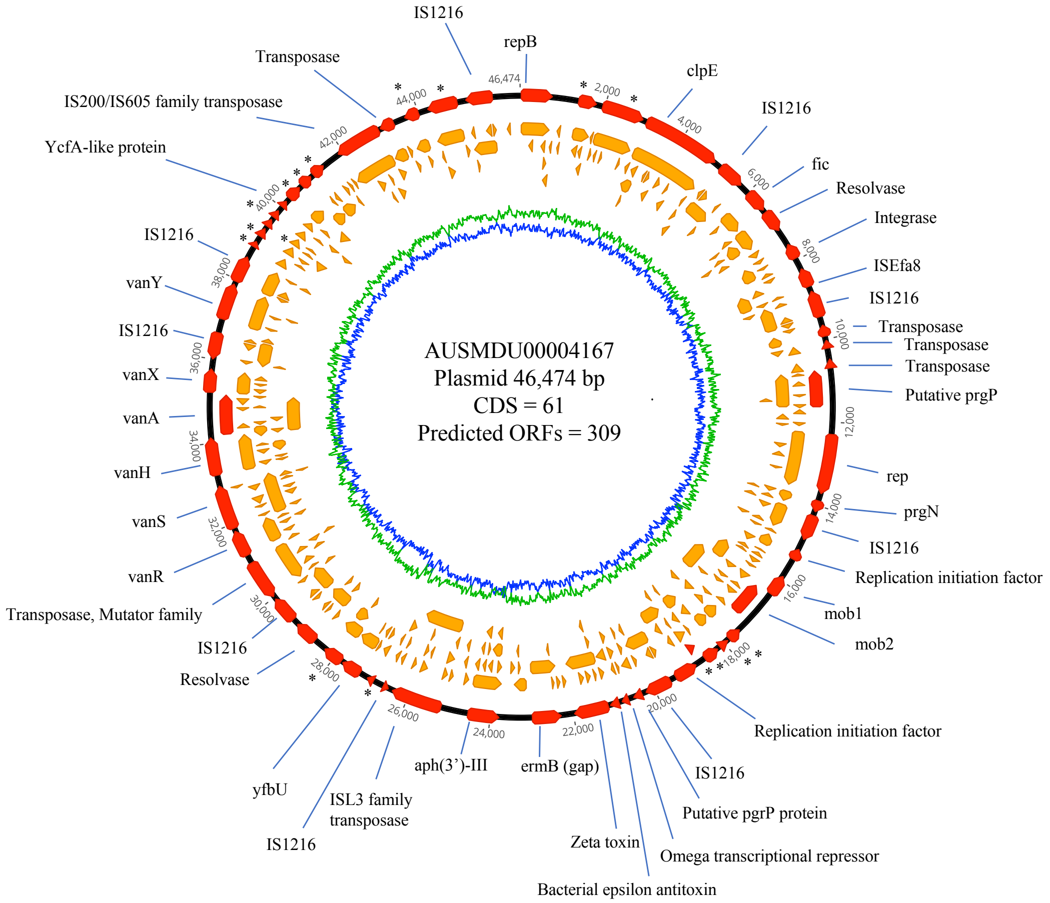


**Legend.** PacBio Single Molecule Real-Time sequencing was performed on a single *vanA*-positive isolate from each of the main STs identified (ST 78, 80, 203, 796, 1421). Plasmids harbouring *vanA* were initially identified using ABRicate (https://github.com/tseemann/abricate), and subsequently annotated using the Enterococcus database (https://github.com/tseemann/prokka/blob/master/db/genus/Enterococcus). Plasmids were then circularized and visualized in Geneious (v.9.1.7, (20)). Open reading frames (ORFs) were also predicted in Geneious, with a minimum of 100 bp required. Annotated genes are shown in red on the outmost circle, and predicted ORFs are orange. GC content is shown in blue, while AT content is shown in green. * represents a hypothetical protein.

**Figure A6.** Homology with closely-related published *vanA*-harbouring plasmids identified in Table S3

AUSMDU00004055 (ST 78) plasmid

AUSMDU00004028 (ST 796) plasmid

AUSMDU00004142 (ST 80) plasmid

pHvH-V24 (Denmark)

AUSMDU00004024 (ST 203) plasmid

pJEG043 (Australia)

Ef_DMG1500501 plasmid 4 (Australia)

AUSMDU00004167 (ST1421) plasmid

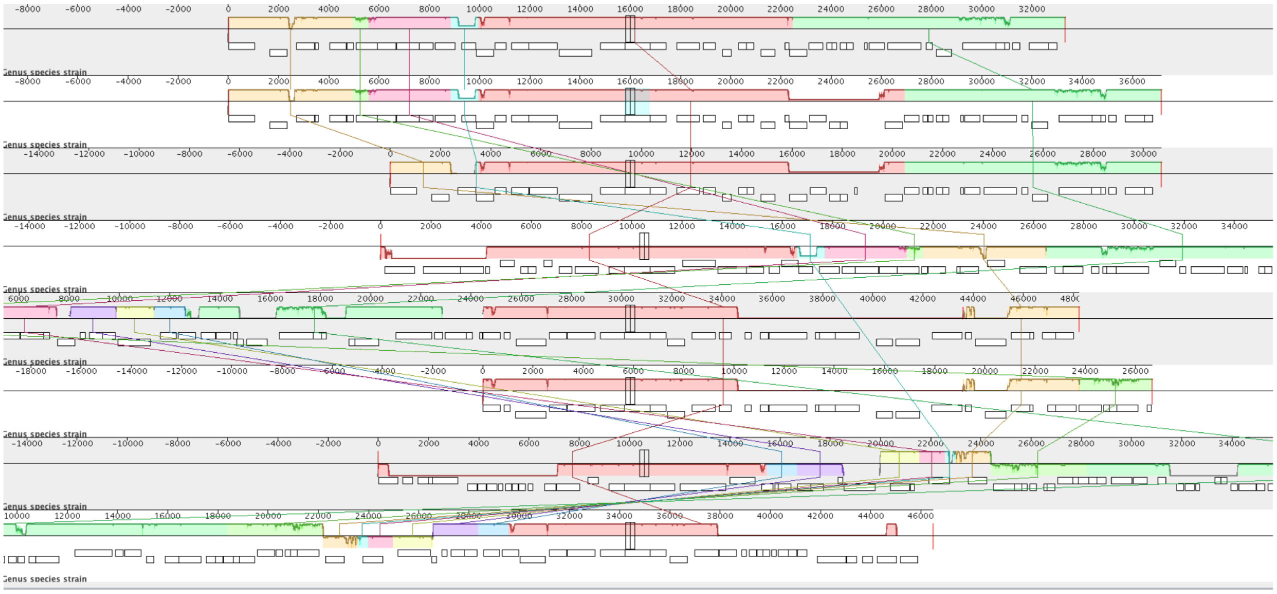

**Legend.** PacBio Single Molecule Real-Time sequencing was performed on a total of five *vanA*-positive isolates, each representing one of the predominant ST types identified (ST 78, 80, 203, 796 and 1421). The *vanA*-harbouring plasmids were identified as described, with annotation done using prokka (v.1.12) based on the Enterococcus database (https://github.com/tseemann/prokka/blob/master/db/genus/Enterococcus). BLASTn (14) was used to identify published plasmids with the greatest sequence homology (**Table A4)**. Published plasmids (from (8, 15, 16)) were likewise annotated with prokka, and alignments were generated using progressiveMauve (build date 2014-12-19). The *vanA* is indicated on each alignment using the black marker.

**Figure A7.** Victorian *E. faecium* in context with published strains

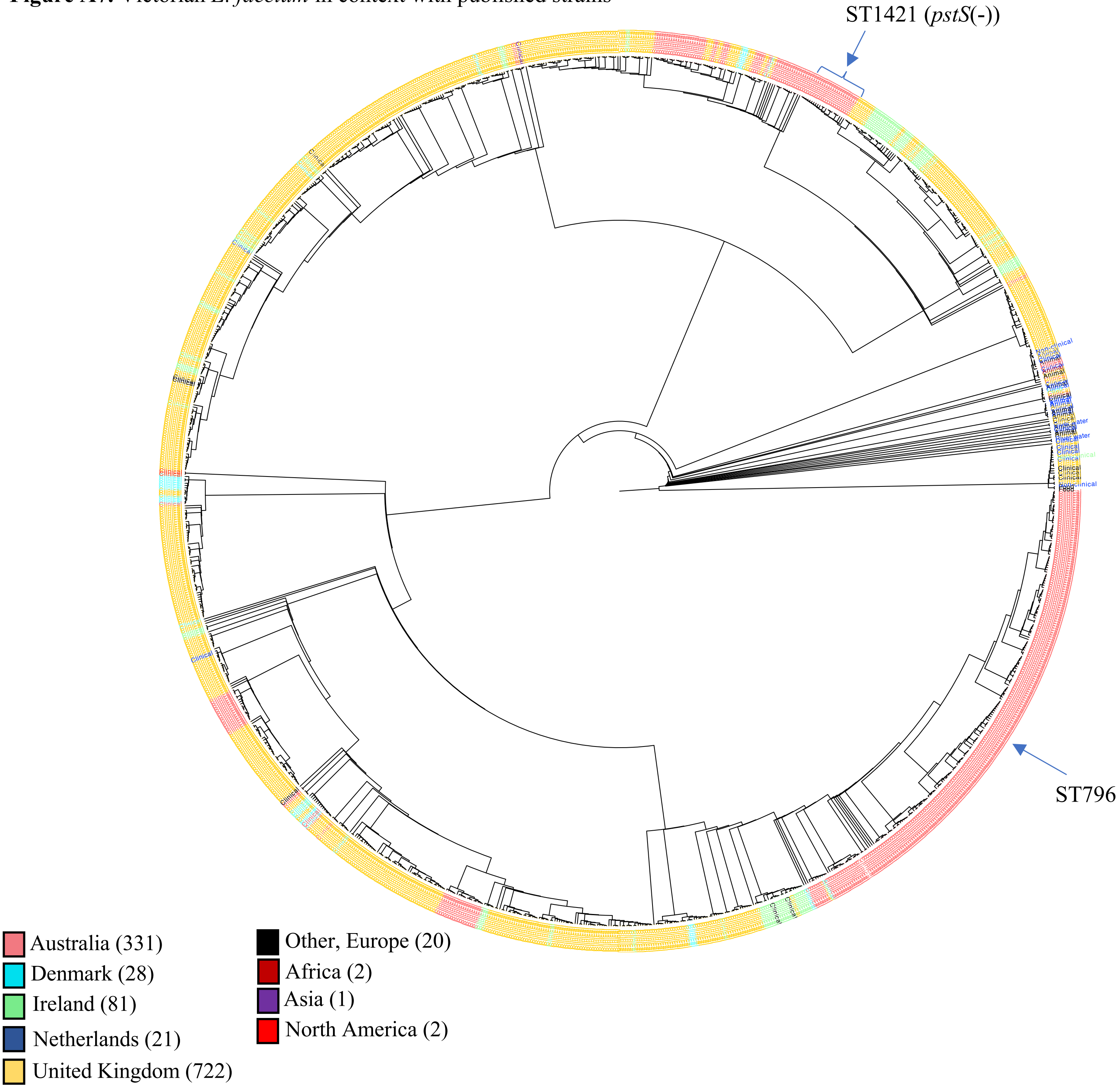


**Legend.** A phylogenetic tree of the 331 Australian isolates, plus 877 isolates from the literature (5-8), based on a concatenated SNP alignment with 111,858 positions. IQ-TREE (v.1.6.1, (9)) was used to generate a maximum likelihood tree, with 20,000 ultra-fast bootstrap replicates. The tree is rooted at the midpoint. For clarity, European countries with fewer than 10 isolates are shown as 'Other, Europe'. The 'United Kingdom' includes isolates that were listed only as United Kingdom, as well as isolates from Wales, Scotland and England. As isolates from the Republic of Ireland and Northern Ireland were not specifically distinguished from one another, these are simply coloured as 'Ireland' above. Where only one isolate was available from a continent, these were coloured by the continent. Isolates were classified as 'Clinical', 'Non-clinical', 'Animal', etc., per previous publications. If an isolate had no previous annotation, but was taken from human blood, it was also classified as 'Clinical' for this analysis. This tree was visualizing using FigTree (v.1.4.2). Branch lengths have been transformed proportionally for easier visualization of tree structure. The most represented ST among *vanA* and *vanB*-VREfm is indicated.
